# The roles of MKI67, GNL2, and MDN1 in Ribosome biogenesis and Transcriptome regulation in the Neuronal Lineage cell line HEK293T

**DOI:** 10.1101/2025.02.13.638155

**Authors:** Shiro Iuchi, Joao A. Paulo

**Affiliations:** Department of Cell Biology, Harvard Medical School, 240 Longwood Ave., Boston, MA, 02115, USA

## Abstract

Ki-67, a protein encoded by the *MKI67* gene, is present in proliferating human tissue cells. It is an important marker for diagnosing cancer. Despite numerous investigations, the function of Ki-67 remains unclear. In this study, we examined the role of Ki-67 and its associated proteins, GNL2 and MDN1, in the HEK293T neuronal lineage cell line using ChIP-seq, RNA-seq, confocal microscopy, and biochemical methods. Ki-67 bound to nearly all regions of the chromosomes through certain consensus nucleotide sequences and was primarily present at the nucleolar periphery, where it interacted with GNL2. These two proteins could recruit MDN1 to the periphery. Depleting Ki-67, GNL2, or MDN1 resulted in characteristic changes in nucleolar protein and chromatin localization. These phenotypes and protein-protein interaction data suggest that Ki-67 regulates the import of nuclear proteins and the export of pre-60S particles. Specifically, Ki-67 retained pre-60S particles within the nucleolus and, when recruiting MDN1, released them along with itself and chromatin. Depleting any one of these proteins decreased the levels of RNAs involved in ribosome biogenesis, with the strongest decrease occurring with MDN1 depletion. MDN1 depletion also decreased transcripts involved in mitochondrial respiration, resulting in strong culture acidification. Conversely, depleting Ki-67 increased the levels of over a thousand different transcripts. The greatest number and strongest increases were of transcripts necessary for neuronal development and activity. These included *PAX5*, *SCG3*, *UNC13A, SLITRK1, CHGB, CHGA, POU4F2, KCNB1, NEK7*, and *SLC7A2*. *SLC7A2* encodes an amino acid transporter that activates mTORC1 with imported amino acids. However, the transcript of NEUROG2, a master regulator that activates downstream genes necessary for forming neuronal structures, did not increase. Taken together, these results suggest that Ki-67, GNL2, and MDN1 promote neuronal cell proliferation by regulating ribosome processing and energy production. Nevertheless, the disappearance of Ki-67 is essential for nervous system development, activity, and survival.

## Introduction

Gerdes and his colleagues initially identified Ki-67 (MKI67) as a marker present only in dividing human tissue cells [1, 2]. This discovery led to the development of a valuable cancer diagnostic tool. Ki-67 exists in two forms. The longer version contains 3,256 amino acids. The shorter version contains 2,896 amino acids, lacking a segment but retaining its main features [1, 3, 4]. Throughout the molecule, Ki-67 contains many regions consisting of charged amino acid residues that form numerous intrinsic disordered regions (IDRs) (Fig 1A).

**Fig 1.**
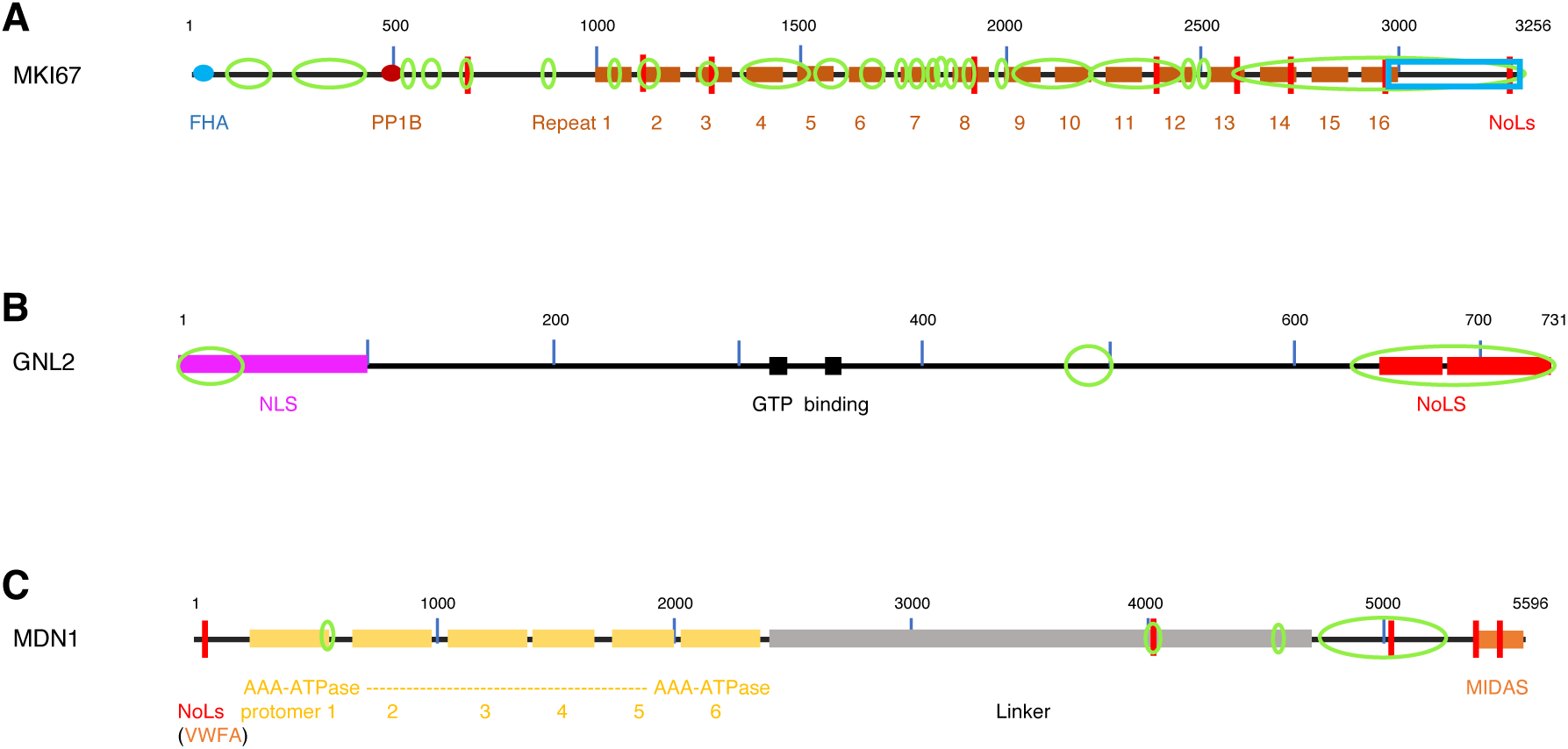
Schematic representations of the primary structures of proteins. **A**. KI-67, **B**. GNL2. **C**. MDN1. The marks indicate the followings: red bar, NoLS; green loop, IDR; blue rectangle, C-terminal DNA-binding region; and pink bar, NLS. Additional structural information is provided directly below each figure.

These IDRs enable Ki-67 to extend within cells [5, 6] and form droplets via liquid-liquid phase separation (LLPS), a process that can be observed in vitro [7]. This propensity appears to explain how Ki-67 interacts with hundreds of proteins [8–10]. In addition to this unique propensity, Ki-67 has domains that specifically interact with ligand proteins. Its N-terminus contains an FHA domain to which NIFK primarily binds during mitosis [11], and its C-terminus contains a 321-residue domain called the LR domain that preferentially binds to AT-rich DNA sequences [12, 13]. Between its N- and C-terminal ends, Ki-67 contains a PP1 motif, to which the PP1γ phosphatase binds [8, 14], and a domain with 16 tandem repeats of 122 residues each [15]. During interphase, Ki-67 binds to chromosomes through its C-terminal LR domain. During mitosis, Ki-67 binds to chromosomes through its LR domain, as well as through the phosphorylation of amino acid residues mostly at the region adjacent to the 16 tandem repeats by CDK1 and other kinases [4, 7, 16, 17]. Consequently, Ki-67 forms a thick perichromosomal layer (PCL) containing nucleolar proteins [8]. The pA-DamID technology has demonstrated that Ki-67 binds to wide regions of chromosomes and that these binding levels and regions change throughout the cell cycle [18]. However, the results of this method unexpectedly show that Ki-67 binds to a limited region of chr2 in randomly growing cells despite its binding to wide regions of chromosomes at most phases of the cell cycle. These underestimated binding regions may result from the indirect identification of Ki-binding sites. Ki-67 undergoes active turnover at the nucleolar periphery during interphase and in the PCL during mitosis [4, 8]. Ki-67 in the PCL acts as a surfactant in keeping sister chromatids apart during mitosis so that they can separate correctly into two sister cells [6]. Before the nuclear membrane reassembles following mitosis, Ki-67 clusters the chromosomes and excludes large cytoplasmic molecules, such as mature ribosomes from the nucleus [19]. Thus, the two roles of Ki-67 in mitosis have been revealed. Conversely, Ki-67’s role in interphase is not well understood, except that Ki-67 is necessary for smooth entry into the S phase and the high expression of genes related to DNA synthesis. Ki-67 maintains low levels of the tumor suppressor protein p21 (CDK1A), allowing cells to enter S phase smoothly. Consistently, cells that do not induce p21 expression, such as HeLa and HEK293T cells, do not experience retard entry to S phase when Ki-67 is depleted [20, 21]. It has been reported that Ki-67 regulates gene expression. However, results among research groups are often inconsistent. For example, one research group found that Ki-67 is necessary for *rRNA* gene expression, while another group reported the opposite [8, 9]. Furthermore, circumstantial evidence supports both types of the results [9, 22–26]. More discrepancies regarding the regulatory role of Ki-67 in gene expression have been reported. However, these discrepancies in the number and types of genes appear to arise from due to differences in organisms, cell types, cell immortalization methodology, and degrees of cell differentiation [9, 18, 27]. The expression of the *MKI67* gene itself has been studied [28]. The DREAM complex, which consists of MuvB and its repressor subunits, binds to both repressive and activating cis elements. This binding mode represses MKI67 transcription during the G0/G1 phases. However, MuvB can replace these subunits with proteins, such as B-MYB, which enables the complex to bind only to the activating cis element of the *MKI67* gene. The *MKI67* transcript levels peak in prophase and mitosis [28–30]. The protein Ki-67 is very low or undetectable during the G0/G1 phase but increase approximately five hours after the mRNA expression increase, ultimately peaking during mitosis. Interestingly, cells experience mitotic DNA damage in the absence of Ki-67 [31], and the LR domain alone can suppress this DNA damage.

GNL2 is a nucleolar GTPase composed of 731 amino acid residues. It enters the nucleolus in two steps, using an NLS at the N-terminus and two NoLSs at the C-terminus (Fig 1B) [32–34]. The *S. cerevisiae* homolog of GNL2, Nog2, binds to the pre-60S complexes to form nucleolar Nog2 particles [35]. These complexes then convert to the Rix1-Rea1 particles in the nucleus. Rea1, the yeast homolog of the human protein MDN1, replaces Nog2 with the export adaptor Nmd3 to form particles that exit the nucleus and enter the cytoplasm. Eventually, the particles mature in the cytoplasm [35–37]. Human pre-60S particles are exported to the cytoplasm via Exportin 5 [38]. Thus, the export mechanism of the pre-60S particles from the nucleus to the cytoplasm is understood; however, the export mechanism from the nucleolus to the nucleus remains unclear. GNL2 interacts with the ribosomal protein RPL23A in human cells [39], so it likely exists in the nucleolus as a subunit of the pre-60S particles, as it does in *S. cerevisiae*. Additionally, GNL2 is abundantly present at the nucleolar periphery, where it interacts with Ki-67 [10]. However, it is not expected to be present in yeast because the organism lacks Ki-67 [40]. Therefore, the role of nucleolar peripheral GNL2 is likely specific to cells of organisms with Ki-67, such as human cells. This type of GNL2 may play a role in transporting substrates in and out through the nucleolar periphery.

Nucleolar contents originate from de novo RNA polymerase I transcription and nucleolar import through the nucleolar periphery although some of them come from the PCL in anaphase [8]. During interphase, RNA polymerase I transcribes the 47S precursor ribosomal RNA (pre-rRNA) from *rRNA* genes located in nucleolar organizing regions (NORs) on the p-arms of acrocentric chromosomes 13, 14, 15, 21, and 22. Meanwhile, the *5S rRNA* is produced on chromosome 1 and imported into the nucleolus [23]. Many other proteins, RNAs, and ribonucleoproteins (RNPs) are also imported into the nucleolus. These proteins, RNAs, and RNPs assemble into pre-60S and pre-40S particles, either directly or after undergoing chemical modification. For example, pre-rRNA undergoes covalent modification by small nucleolar RNAs (snoRNAs), which alters its structure to facilitate cleavage into smaller rRNA species [41]. Then, these RNAs are cleaved and mature into 18S, 5.8S, and 28S rRNAs. Next, the resulting RNAs and proteins assemble into pre-60S particles. These proteins include RPLs, as well as the RIX1 complex (PELP1, NOL9, TEX10, WDR18, and SENP3), GNL2, and MDN1 [36, 41–45]. MDN1, a large AAA-ATPase consisting of 5,596 amino acid residues (Fig. 1C), is recruited by SUMOylated PELP1 [46] and plays a crucial role in processing the pre-60S particles. Current knowledge of particle processing and the role of each protein largely depends on advanced research results from *S. cerevisiae* [35, 36]. In *S. cerevisiae*, the Rea1 (MDN1) migrates with the pre-60S particle from the nucleolus to the nucleus through the cytoplasm, where it catalyzes the final processing steps [36, 37]. In human cells, MDN1 interacts with GNLs and Ki-67 at the nucleolar periphery. This additional step in pre-60S particle processing suggests that the MGM complex plays an important role in processing at the human nucleolar peripheral boundary.

Inside cells, proteins often serve as vital components of large macromolecular complexes, such as ribosomes, spliceosomes, and the CCR4-NOT complex. These complexes carry out specialized biosynthetic or degradative functions [36, 47–49]. However, we demonstrated that these complexes are interconnected in a larger metabolic network called the RNAmetasome. Furthermore, we discovered that Ki-67 (MKI67), GNL2, and MDN1 occupy a central position in this network by forming a complex called the MGM at the nucleolar periphery [10]. This finding prompted us to investigate the functions of the MGM complex and its proteins in HEK293T neuronal lineage cells. In this report, we reveal details about Ki-67 binding to chromosomes, MGM complex formation, and the complex’s role in ribosome maturation. We also reveal that these proteins regulate genomic transcript levels. MDN1 plays a crucial role in coordinating RNA levels for ribosome biogenesis and mitochondrial respiration. Ki-67 silences the expression of numerous neuron-specific genes, making its absence essential for neuronal differentiation, activity, and survival. Additionally, we discuss Ki-67’s involvement in the formation of nucleolus-associated domains (NADs).

## Materials and Methods

### Cell culture

The HEK293T cell line (ATCC CRL-3216) was cultivated in DMEM (Thermo Fisher Scientific) supplemented with 10% fetal bovine serum (HyClone). The cultivation and the handling procedures of this cell line have been approved by the Harvard University Committee on Microbiological Safety (ID: 20-155). All experiments were conducted in accordance with the NIH rDNA Guidelines and the Federal Occupational Safety and Health Administration Bloodborne Pathogen Standard Guidelines. HEK293T cells overexpressing the MKI67 (Ki67) transcripts were established by selecting cells resistant to 0.5 µg/ml puromycin after transfecting the cells with the EGFP-Ki67 plasmid (HEK293T/EGFP-Ki67), and the control cells were established using the EGFP plasmid (HEK293T/EGFP). The EGFP-Ki67 plasmid was constructed by removing the mCherry gene from the EGFP-Ki67-mCherry plasmid [6] with BamHI-HF and NotI-HF, followed by filling and ligation (S1A Table). Similarly, the EGFP plasmid was constructed, by digesting the original plasmid with BsrGI-HF and NotI-HF to remove the Ki67 and mCherry genes.

### Cloning of the *GNL2* cDNA and its truncation

Total RNA was purified from HEK293T cultures, and the first-strand cDNA was synthesized from 600 ng of the total RNA using the ProtoScript II Kit (NEB, E6560S). Subsequently, the cDNA encoding GNL2 (Q13823) was amplified using Q5 Hot Start High-Fidelity DNA Polymerase (NEB, M0494S) and cloned into the ECORI-SCAII site of pEGFP-C2, yielding pGNL2-G7 (S1A Table). This plasmid encodes GNL2 tagged with two tags with an N-terminal EGFP tag and a C-terminal 6x His tag. The 6x His tag was then removed, yielding pGNL2-#1. Subsequently, the two plasmids encoding full-length GNL2 were differentially truncated using the Q5 Site-Directed Mutagenesis Kit (NEB). Similarly, pGNL2-G7 and the fragments were ligated to pGEX-5X-1 at the ECORI-XhoI site to yield bacterial expression plasmids. All oligonucleotides used for cloning, truncation, and related plasmid construction were purchased from IDT (S1A Table).

### Knockdown (KD) of genes

The HEK293T culture was transfected with gene-specific DsiRNA (Dicer-substrate small interfering RNA for gene knockdown, IDT) at 40 nM concentration according to the reverse transfection protocol of Lipofectamine RNAiMAX (ThermoFisher) to knockdown the *MKI67, GNL2, MDN1, YTHDC2, CNOT1, G3BP1, G3BP2*, and *FMR1* genes. Similarly, the HEK293T culture was transfected with a universal negative control RNA (NC1) to create the control KD cultures. For an experiment, a set of the transfected cultures was harvested after an incubation period ranging from 65 to 70 h to avoid harvesting the saturated cultures. The cells were predominantly cultured in 12-well plates, but they were also cultured in 35-mm Matsunami glass-bottom dishes for cytological work. The DsiRNAs used in this study were purchased from IDT (S2 Table).

### Western blot for analysis of the cellular interaction between GNL2 and endogenous Ki-67 or MDN1, as well as for analysis of gene knockdown

To investigate the native protein-protein interactions, HEK293T cells grown in a 10-cm dish were transfected with 10 µg of a plasmid encoding either full-length GNL2 or its truncated protein using X-tremeGENE 9 DNA transfection reagent (Sigma). The cultures were cultured for two days. Nuclear extracts were prepared using the nuclear extraction buffers supplemented with 10 µM BTZ (Bortezomib, AdipoGen LIFE SCIENCES, AG-CR1-3602), after the cells were exposed to a hypotonic buffer as previously described [10]. The nuclear extracts (300 µl) were precleared with a mixture of 20 µl of protein A magnetic beads and protein G magnetic beads for 60 min. The precleared nuclear extracts were then incubated with 3 µg of either rabbit anti-GFP antibody or mouse anti-6xHis antibody for 60 min. They were then incubated for 70 min after adding 20 µl of a mixture of protein A and protein G magnetic beads. The beads were washed 4 times with 300 µl of the nuclear extraction buffer containing 10 µM BTZ, suspended in 30 µl of 1.5x loading buffer. and finally heated at 100C for 5 min. Subsequently, 10 µl of the supernatant was applied to one well of 4–20% Mini-PROTEAN TGX gels (BIO-RAD, 4561096) for electrophoresis. The immunoprecipitants separated on the gel were transferred to a PVDF membrane. The proteins on the membrane were then detected using antibodies at the concentrations described (S3 Table). To investigate the effect of gene knockdown, KD cultures were trypsinized and collected in 1.5-ml centrifuge tubes and washed twice with PBS/1x Complete EDTA-free protease inhibitor cocktail (Roche). The cell pellets were resuspended in 30 µl of hypotonic lysis buffer, kept on ice for 20 min and subjected to two freeze-thaw cycles. The resulting cell lysates were then used as cell extracts. After estimating the protein concentration using the Micro BCA kit (Fisher Scientific), 50 µg of protein was loaded into one well of 4–20% Mini-PROTEAN TGX gels (BIO-RAD 4561094). The loaded proteins were separated by PAGE and then subjected to Western blot analysis using the antibodies (S3 Table).

### Far western for in vitro protein-protein interaction analysis

A prey sample was loaded and separated by SDS-PAGE, then transferred to a PVDF membrane. The membrane was immersed in 20 mL of a 6 M guanidine HCl solution and incubated for 10 minutes. Then, the membrane was transferred to a solution with half the concentration of guanidine HCl [50] and incubated for 10 minutes. This transfer followed by a 10-minute incubation was repeated six times to gradually decrease the guanidine concentration to less than 100 mM. The membrane was washed twice with PBS, blocked with LI-COR blocking solution for one hour, and incubated with a probe for two hours at room temperature (RT). Purified #20 or #21 was used at the concentration (1.1 x 10⁶ LI-COR units) to examine their binding affinity for the prey sample. After washing the membrane in PBS/0.1% Tween 20, it was subjected to double protein staining with a combination of primary mouse and rabbit antibodies, followed by visualization with secondary antibodies (S3 Table).

### Confocal microscopy

The cultures grown on Matsunami glass-bottom dishes were rinsed once with serum- and antibiotic-free DMEM and then fixed with 4% paraformaldehyde/PBS at room temperature for 10 min. Fixation was terminated with 3x PBS, followed by replacement with 1x PBS. The fixed cultures were then dehydrated sequentially with 50%, 75%, 95%, and 100% ethanol and then air-dried. Subsequently, the cells were permeabilized with 0.2% Triton X-100/PBS for 10 min and then blocked with 1x PBS/5% BSA/0.1% IGEPAL CA630 for at least 5 min. The samples were typically double-stained with mouse and rabbit primary antibodies in the blocking buffer at 4C for 16-18 h and then washed four times with the blocking buffer (S3 Table). For the control staining, the blocking buffer without antibodies was used. They were further stained with two different goat secondary antibodies conjugated to a dye in the blocking buffer for 1.5 h at room temperature. Then, they were washed three times and rinsed once with water and eventually air-dried. The samples were treated with DAPI-containing Vectashield (Vector) and examined with a confocal microscope, Lucille spinning disk & FR, at the CITE at Harvard Medical School. Immunofluorescence images were acquired by capturing the emission from the excitation of the Alexa Fluor 555 (red) and Alexa Fluor 488 (green) dyes, as well as the DAPI emission. Z-plane images of a cell (or culture field) were taken every 0.2 µm. Of these images, the one cut through the center of the nucleolus was used for comparison between images of cells that were cultured under different conditions, unless otherwise noted. The acquired images were visualized by subtracting nonspecific signals derived from the control and analyzed using Fiji [51].

### Probe preparation

The E. coli strain (NEB C3013) was transformed with plasmid #20, #21, or #22, and grown in 30 ml of LB containing 100 µg/mL100-µg per ml carbenicillin to an early log phase and then induced with 0.5 mM IPTG for 4 h to produce the encoded protein from the plasmid. The cells were pelleted by centrifugation and suspended in 0.3 ml of 150 mM NaCl/25 mM Na-phosphate buffer (p8.0)/2x Complete (EDTA-free)/1mM PMSF/0.5-mg/ml lysozyme, subjected to freeze-thawing once. Then, the cells were sonicated twice at 50% output power for 5 seconds using a Branson sonicator. The extract was centrifuged at 14000 rpm for 10 min. The supernatant was mixed with Triton X-100 to reach a concentration of 0.05%. Two hundred and fifty µl of the extract were mixed with an equal volume equilibration buffer (50 mM sodium phosphate/0.3 M NaCl/10 mM imidazole/0.05% Tween 20, pH 8.0). Then, the sample mixture was applied to 50 µl of washed and diluted twofold Ni-NTA magnetic agarose (Thermo Scientific 78605). Subsequently, this agarose-extract mixture was rotated at 4C for 1 h, washed four times with 250 µl of wash buffer (50 mM sodium phosphate/0.3 M NaCl/15 mM imidazole/0.05% Tween 20, pH 8.0), and then the bound probe was eluted once with 50 µl of elution buffer A (50 mM sodium phosphate/0.3 M NaCl/0.1 M imidazole, pH 8.0). The remaining probe on the beads was then eluted twice with 50 µl of elution buffer B (50 mM sodium phosphate/0.3 M NaCl/0.3 M imidazole, pH 8.0). The last two elutes were combined and used as the probe for the Western blot.

### Preparation of protein samples from KD cultures

The cultures grown in 12-well plates were rinsed once with PBS, after which they were trypsinized with 500 µL of a 0.025% trypsin/0.26 mM EDTA solution for five minutes. Trypsinization was terminated by adding 500 µL of fresh medium. The cell suspensions were transferred to 1.5-ml centrifuge tubes and washed twice with 500 µL of PBS containing an EDTA-free protease inhibitor cocktail at 4 °C. The cells were resuspended in 30 µL of a hypotonic lysis buffer containing 10 mM KCl, 10 mM HEPES (pH 7.4), 0.05% IGEPAL CA-630, 0.2 mM sodium orthovanadate, 2X EDTA-free protease inhibitor cocktail, 2 µL/mL benzonase, and 2 mM MgCl₂. The suspensions were placed on ice for 20 minutes and then subjected to freeze-thawing twice. Cell lysates were obtained.

### Protein samples for MS analysis

Each of four protein samples (approximately 100 µg protein) prepared from duplicate KD-MKI67 and duplicate KD-CTL cultures was mixed with 10-volumes of cold 10%TCA/acetone, vortexed, and placed in a −20C freezer overnight. Precipitated proteins were collected by centrifugation at 15,000 rpm for 10 min at 4C and resuspended in 10-volumes of cold acetone of the original sample, vortexed, placed on ice for 10 min, and centrifuged at 15,000 rpm for 5 min at 4C. Proteins were washed with the 10 volumes of cold methanol of the original sample and air dried for 15 min. The samples were digested in 100µL of 100 mM EPPS, pH 8.5 and at 37°C with trypsin at a 100:1 protein-to-protease ratio overnight. The samples were desalted via StageTip, dried via vacuum centrifugation, and reconstituted in 5% acetonitrile, 5% formic acid for LC-MS/MS processing.

### Mass spectrometric data collection

Mass spectrometry data were collected using a Exploris 480 mass spectrometer (Thermo Fisher Scientific, San Jose, CA) coupled with a Proxeon 1200 Liquid Chromatograph (Thermo Fisher Scientific). Peptides were separated on a 100 μm inner diameter microcapillary column packed with ∼25 cm of Accucore C18 resin (2.6 μm, 150 Å, Thermo Fisher Scientific). We loaded ∼1 μg onto the column. Peptides were separated using a 90min gradient of 3 to 22% acetonitrile in 0.125% formic acid with a flow rate of 350 nL/min. The scan sequence began with an Orbitrap MS^1^ spectrum with the following parameters: resolution 60,000, scan range 350−1350 Th, automatic gain control (AGC) target “standard”, maximum injection time “auto”, RF lens setting 50%, and centroid spectrum data type. We selected the top twenty precursors for MS^2^ analysis which consisted of HCD high-energy collision dissociation with the following parameters: resolution 15,000, AGC was set at “standard”, maximum injection time “auto”, isolation window 1.2 Th, normalized collision energy (NCE) 28, and centroid spectrum data type. In addition, unassigned and singly charged species were excluded from MS^2^ analysis and dynamic exclusion was set to 90 s. Each sample was run twice, once with a FAIMS compensation voltage (CV) set of −40/−60/−80 V, and the second with a CV set of −30/−50/−70 V. A 1 s TopSpeed cycle was used for each CV.

### Mass spectrometric data analysis

Mass spectra were processed using a Comet-based in-house software pipeline. MS spectra were converted to mzXML using a modified version of ReAdW.exe. Database searching included all entries from the humanUniProt database (downloaded September 2023), which was concatenated with a reverse database composed of all protein sequences in reversed order.

Searches were performed using a 50-ppm precursor ion tolerance. Product ion tolerance was set to 0.03 Th. Oxidation of methionine residues (+15.9949 Da) was set as a variable modification. Peptide spectral matches (PSMs) were altered to a 1% FDR [52, 53]. PSM filtering was performed using a linear discriminant analysis, as described previously [54], while considering the following parameters: XCorr, ΔCn, missed cleavages, peptide length, charge state, and precursor mass accuracy. Peptide-spectral matches were identified, quantified, and collapsed to a 1% FDR and then further collapsed to a final protein-level FDR of 1%. Furthermore, protein assembly was guided by principles of parsimony to produce the smallest set of proteins necessary to account for all observed peptides. Spectral counts were then extracted, and the data were subsequently analyzed.

### ChIP-seq

A HEK293T culture was cultivated to a density of approximately 5 x 10⁷ cells in a 15-cm dish. The cells were washed once with cold PBS and then crosslinked with 300 mM EGS (Thermo Fisher Scientific) in PBS for 30 minutes. Subsequently, a second crosslinking step was performed, employing 1% formaldehyde. The crosslinking reaction was terminated by a 5-minute incubation with 125 mM glycine. Thereafter, the cells were washed and suspended in 3 ml of PBS and transferred to a centrifuge. This collection procedure was repeated to thoroughly collect the cells. The crosslinked cells were once washed with PBS, suspended in 400 µl of lysis buffer (50 mM HEPES-KOH pH8.0/140 mM NaCl/1 mM EDTA pH8/1% Triton X-100/0.1% Sodium Deoxycholate/0.5% SDS/10 µM BTZ). The cells were then placed on ice for 10 min and subsequently sonicated for 10 seconds at one-minute intervals on ice at 30% output power using a Branson sonicator. This is the general procedure for preparation of ChIP-seq samples. However, two distinct operating conditions were employed in this study: 1) Eleven cycles of sonication were performed on 5 x 10⁷ cells that had been treated with formaldehyde for 10 minutes; 2) Thirteen cycles of sonication were performed on 10 x 10⁷ cells that had been treated with formaldehyde for 12 minutes. The prepared extract was diluted fivefold with dilution buffer (lysis buffer lacking SDS) and centrifuged at 14,000 rpm for 10 minutes. The resulting supernatant was precleared with a mixture of protein A and protein G magnetic beads (NEB) at 4°C for one hour. Thereafter, the supernatant was divided into four parts. Each supernatant was supplemented with 4 µg of rabbit anti-Ki-67, rabbit anti-GNL2, rabbit anti-MDN1, or rabbit anti-GFP (serving as the control) and 60 µl of the protein A and protein G magnetic beads mixture. Then, they were then rotated at 4°C for 16 hours for immunoprecipitation. The beads were washed three times with 500 µl of a high-salt buffe (50mM HEPES-KOH pH 8/500 mM NaCl/1 mM EDTA/1% Triton100/0.1% SDS/0.1% Sodium Deoxycholate), washed once with 500 µL of LiCl high salt buffer (50mM HEPES-KOH pH 8/250 mM LiCl/1 mM EDTA/1% Triton100/0.1% Sodium Deoxycholate), and then suspended in 100 µl of elution buffer (50 mM Tris HCl/10 mM EDTA/1% SDS). The crosslinked DNA-protein samples were eluted by incubating the beads at 65C for 5 min, and the crosslink of the sample was removed by incubation at 65C overnight in the presence of 1 µl RNase A (20 mg/ml, NEB). Then, the proteins in the sample were digested with 2 µl proteinase K (800 u/ ml, NEB) for 1 h at 55C. Thereafter, the DNA fragments of the sample were purified using NEB PCR & DNA Cleanup Kit and dissolved in TE buffer. The concentration of the DNA fragments was determined using a Qubit fluorometer (Invitrogen). From these collected DNA fragments (20 µg), a sequencing library was prepared using the NEBNext Ultra II DNA Library Prep Kit for Illumina (NEB E7645). The library was treated with NEBNext Sample Purification Beads (E71044S) twice during the procedure, once to remove the adapter and once to clean up the end product. The High Sensitivity D1000 ScreenTape analysis showed that the end product contained DNA fragments averaging 400 bp in length with no evidence of the adapter contamination. This end-product library was subsequently sequenced to obtain 50 bp paired-end reads on an Illumina NextSeq 2000 sequencer at the Biopolymers Facilities at Harvard Medical School. The first and the second Ki-67 ChIP-seq yielded 125x106 and 240x106 reads, respectively. This is consistent with the expectation resulting from the fact that these ChIP-seqs were initiated with the twofold different numbers of cells.

### Statistical analysis of ChIP-seq

The quality of the sequence data was evaluated using FASTQC v0.11.9 (see FASTQC in the Data and materials availability for reference). Reads were filtered and trimmed with Atropos v1.1.31 [55] to retain high quality reads, which were then mapped to the human reference genome hg38 using Bowtie2 v2.2.5 [56]. Multi-mapping reads and duplicates were removed followed by peak calling with MACS2 v2.2.7.1 [57]. Likely peak artifacts were filtered out using the ENCODE blacklist [58]. ChIP-seq data quality was assessed using standard ENCODE metrics, including total reads, alignment rates, fraction of reads in peaks, PCR bottleneck coefficients and the non-redundant fraction [59]. Data was visualized using IGV [60]. DeepTools v3.0.2 [61] was used to assess coverage and the reproducibility of peaks across replicates. ChIPseeker v1.32.0 [62] was used to annotate ChIP-seq peaks to their closest transcriptional start site and plot binding profiles. HOMER v4.11.1 [63] was used to identify enriched transcription factor binding motifs. Analyses in R were done using R v4.2.1. The MAFFT program in Lasergene 17 was used to align DNA sequences in the centromeric regions to find the long Ki-67 consensus binding sequences.

### RNA-seq

Triplicate cultures of KD-MKI67 (knockdown for the *MKI67* gene) and the KD-CTL (knockdown for control) were cultivated for 70 h after being transfected with 40 nM DsiRNA specific for the MKI67 gene and 40 nM negative control DsiRNA (NC1), respectively. In a separate experiment, triplicate cultures of KD-GNL2, KD-MDN1, and the KD-CTL were cultivated in the same manner after being transfected with 40 nM DsiRNA specific for the *GNL2* and *MDN1* genes, and 40 nM NC1, respectively. The cultures were harvested by trypsinization, and the total RNA was extracted using the Monarch Total RNA Miniprep Kit (NEB T2010S). The total RNA (1000 ng) was subjected to ribosomal RNA depletion using the NEBNext rRNA Depletion Kit v2 (E7400S), and the RNA sample was purified using NEBNext Sample Purification Beads (E7767S). The samples were fragmented by heating at 94C for 10 min.

Approximately 200 bp RNA was used to synthesize indexed cDNA using the NEBNext Ultra Directional RNA Library Prep Kit for Illumina (E7765) and NEBNext Multiple Oligos for Illumina (E7335). The NEBNext RNA Sample Purification Beads (E7767S) were used for both the size selection of the adaptor-ligated DNA and cleanup of the endpoint DNA sample. High Sensitivity D1000 ScreenTape test showed that the resulting cDNA libraries contained an average DNA size of 300 bp and lacked the index primers and adapters. The library was sequenced to obtain 50 bp paired-end products on an Illumina NextSeq 500 system at the Biopolymers Facilities at Harvard Medical School. We analyzed the results using software in the RStudio software (R/4.2.1) on o2, a platform for Linux-based high-performance computing platform managed by the Research Computing Group at Harvard Medical School. The following software was used: Rhisat2 VN2.2.1 [64] to align RNA-seq reads to the reference genome (GRCh38_snp_tran/genome_snp_tran), featureCount in Rsubread V2.12.2 [65]to quantify the number of reads aligned to each gene, and DESeq2 v 1.46.0 [66] to analyze the count matrix for differential gene expression. We used Rsamtools V2.22.0 [67] to sort and index BAM files for visualization in IGV [60].

### qPCR

qPCR was performed in two steps: first, first strand cDNA synthesis and second, amplification of the target cDNA. The first step was performed with the NEB LunaScript RT SuperMix Kit (E3010), and the second step was performed with the NEB Luna Universal qPCR Master Mix (M3003) using Thermo fisher Appliedbiosystems QuantStudio 7 Flex. The two-step cycling conditions recommended by NEB were employed for amplification of target and RPS8 mRNAs. The RPS8 was consistently expressed across a series of culture manipulations according to our RNA-seq data. Subsequently, it was used as an internal reference mRNA to calculate ΛΛCt of target mRNAs. The Ki-67 complementation test was performed using HEK293T/EGFP-Ki67 cells (HEK293T cells stably transfected with the EGFP-MKI67 gene) and control HEK293T/EGFP cells (HEK293T cells stably transfected with the empty vector). The primers used for qPCR are listed in S1B Table.

## Results

### HEK293T is a human neuronal lineage cell line

The HEK293T parent cell line, HEK293, was originally established by transforming cells derived from human embryonic kidney cells with fragmented adenovirus type 5, Ad5 DNA [68]. For this reason, HEK293 was traditionally considered a kidney cell line. However, subsequent research showed that Ad5 preferentially transforms neuronal cells, suggesting that HEK293 has a neuroectodermal origin [69]. This idea has not yet been universally accepted. To clarify the cell origin of the HEK293T cells used in this study, we revisited and analyzed our previous mass spectrometry data using STRING within the Cytoscape platform [50, 70]. Out of the 7,811 proteins identified, 7,570 were matched with high confidence by the database. Tissue enrichment analysis revealed that these proteins are predominantly expressed in the human nervous system, surpassing their expression in kidney or adrenal tissues (S1 Fig). However, the dataset lacked the transcription factors NEUROG2, which are essential for neurogenesis, including the formation of dendrites and axons [50, 71]. Based on these findings, we conclude that the HEK293T cells originated from the neuronal lineage of the embryonic kidney. These immortalized cells most likely originated from differentiated neurons and lost or regressed NEUROG2 expression, preventing the formation of neuronal structures. Consequently, these cells retain neuronal protein expression while proliferating like neural primordial/progenitor cells (NPCs). HEK293T cells, which were originally called 293/tsA1609neo, were established through the second-round immortalization using temperature-sensitive SV40 T-antigen DNA. Since Ad5 and SV40 T-antigen DNA inactivate p53 and pRB through different mechanisms [68, 72–74], the resulting HEK293T cells do not increase p21 expression. Consequently, the cells also do not exhibit two Ki-67 depletion phenotypes: delayed entry into S phase and reduced levels of DNA synthesis transcripts [21].

### Overall features of Ki-67 binding to the chromosomes

The chromosome-MGM complex model was previously established based on the results that reported the binding of Ki-67 to chromosomes [13, 18]. The binding feature of Ki-67 is the most fundamental factor of the model, and it may differ from cell type to cell type. Therefore, we performed Ki-67 ChIP-seq on biological duplicate samples and analyzed the datasets using Integrative Genomic Viewer (IGV). The analysis revealed several binding features of Ki-67.

First, the BigWig track showed that Ki-67 binds to nearly all regions of the chromosomes, excluding regions where input reads were not assigned (Fig 2A and 2B; S2 Fig).

**Fig 2.**
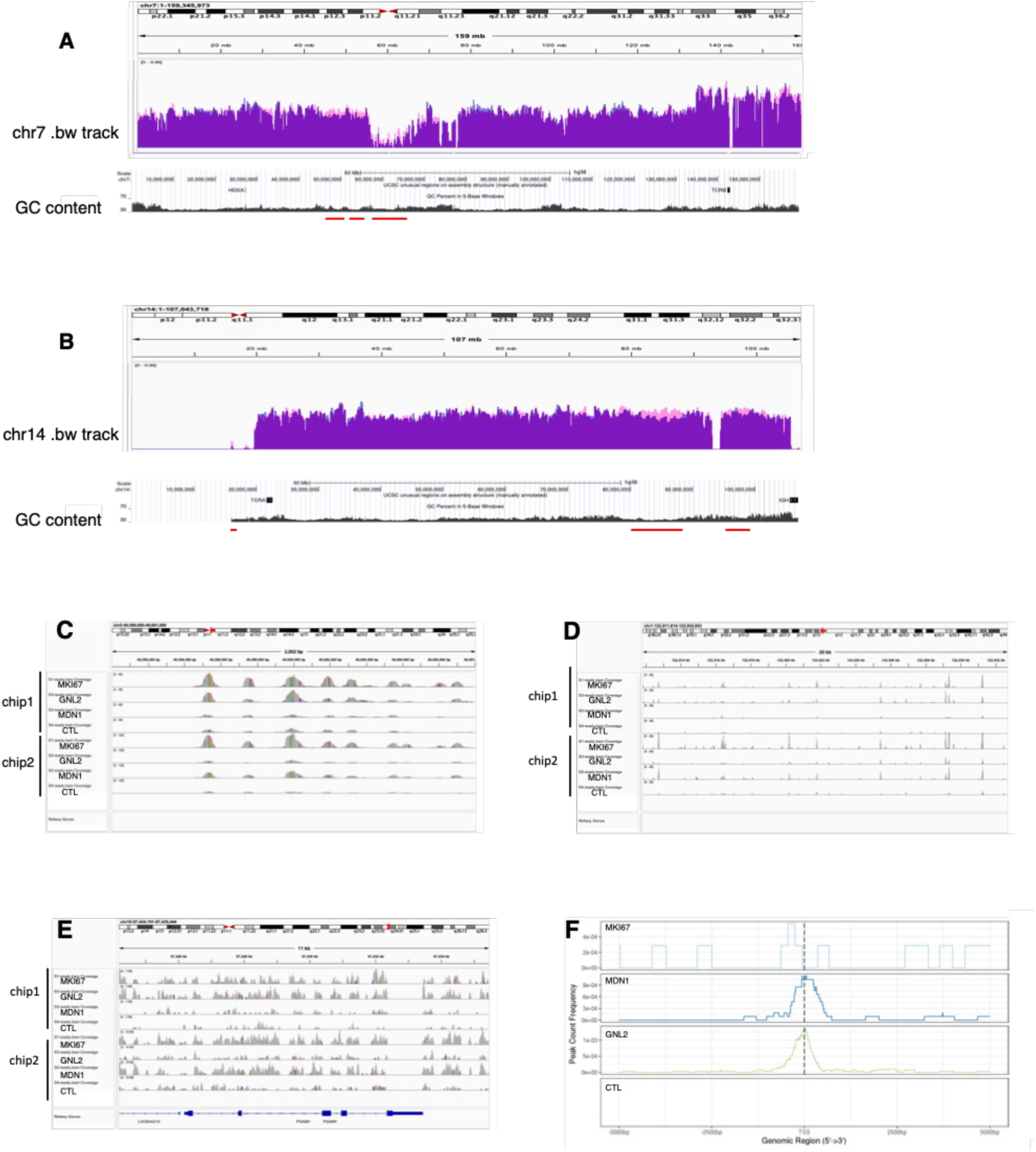
Binding of Ki-67, GNL2, and MDN1 to chromosomes. **A**. Presentation of the bigWig file for the Ki-67 ChIP-seq read track and its input read track on chromosome 7. The two normalized tracks are merged for comparison. Pink, Ki-67 ChIP-seq; blue, input. The two tracks overlap at nearly all regions (violet). Regions to which Ki-67 bound more strongly were located in some AT-rich sequence regions as highlighted by the red lines. **B**. BigWig tracks on chr14 are presented in a similar manner to panel A. **C-E**. A comparison of two independent ChIP-seq BAM files: chip1 (the initial ChIP-seq) and chip2 (the second ChIP-seq). The chromosomal locations of the Ki-67 (C), GNL2 (D), and MDN1 (E) BAM files are indicated by red bars at the top of each file. **F**. The binding of MKI67, MDN1, and GNL2 at promoter regions.

These bindings were consistent between the two datasets (S2 Fig). The BAM tracks of the ChIP-seq analysis provided more detail on the consistency of Ki-67 binding between the datasets (Fig 2C–2E). Second, the results showed that Ki-67 bound preferentially to AT-rich (i.e., GC-poor) regions, though not to all of them (Fig 2A, 2B). This Ki-67 binding preference feature was also observed at the nucleotide sequence level (S3A, S3B Fig), suggesting that factors other than AT-rich sequences are involved in Ki-67 binding. Third, Ki-67 bound to promoter regions, however, this protein does not appear to be a typical transcription factor (Fig 2F) because it also bound to sites adjacent to the transcription start site (TSS). These three observations present Ki-67 binding features in interphase cells because the antibody we used in this study did not virtually detect mitotic Ki-67 of the randomly proliferating cultures (S4 Fig).

### Ki-67 Nucleotide Sequence Specificity

HOMER analysis across the Ki-67 ChIP-seq datasets identified 15 octanucleotide candidates with p-values below 1E-9 (S4 Table). However, due to the unusually long DNA-binding domain of Ki-67, the actual Ki-67-binding DNA sequences may be longer than these octanucleotides. Consequently, we searched for longer Ki-67 binding sites in the centromeric regions (chr1, 2, 3, 4, 5, 7, and 13), where nearly all Ki-67 binding sites existed as single peaks, as shown in Fig 2C, 2D. We searched the centromeric regions instead of the noncentromeric regions to avoid picking up incorrect sequences caused by continuous Ki-binding, which typically occurs in the noncentromeric regions. Each centromeric peak spanned 50–120 bp of DNA (Fig 3A–F).

**Fig 3.**
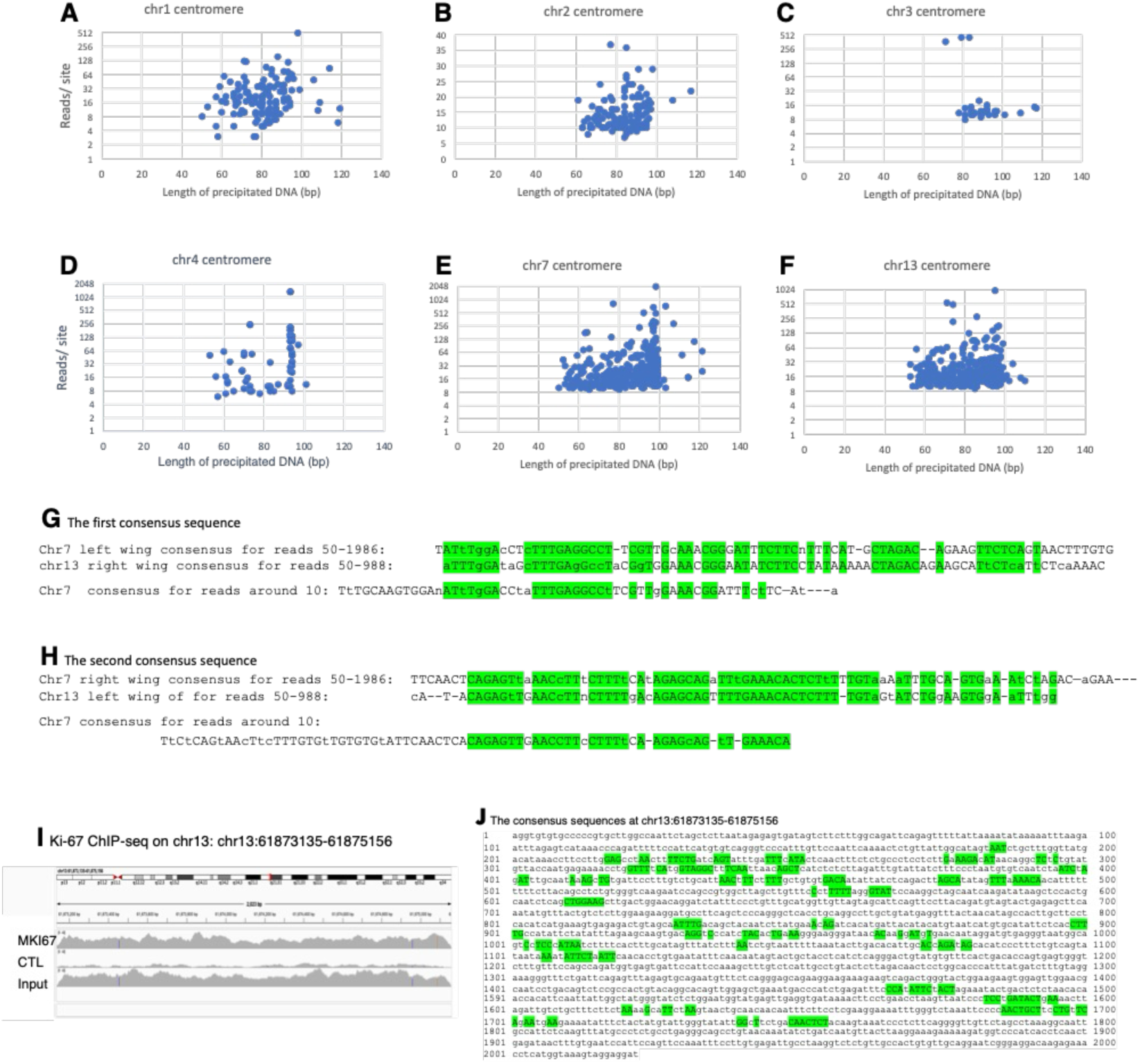
Details of Ki-67 binding to chromosomes. The data presented in this figure derive from chip2 results. **A–F**. Binding of MKI67 to the centromeric regions of chr1, 2, 3, 4, 7, and 13. The x-axis plots the nucleotide sequence length (bp) occupied by Ki-67, and the y-axis plots the number of reads exceeding eight in a peak area. **G**. The first long consensus sequences to which Ki-67 binds. The first and second rows display the consensus sequences on chromosomes 7 and 13, respectively. The third row shows a short consensus sequence found in the chr7 centromere. **H**. The second long consensus sequence to which Ki-67 binds. The first and second rows display the consensus sequences on chromosomes 7 and 13. The third row displays a short consensus sequence found in the Chr7 centromere. **I**. The chr13:61873135-61875156 region to which Ki-67 binds most strongly. **J**. Consensus sequences present on the + or - strand of the chr13:61873135-61875156 region. The consensus nucleotide sequences highlighted in green are parts of two consensus sequences, and these sequences are conserved with the spaces between them.

Interestingly, the peak intensity greatly varied among chromosomes. The greatest peak, i.e., the strongest binding, occurred on chr7, followed by chr13, and then the others. This suggests that chr7 has the most highly qualified Ki-67-binding DNA sequences. Aligning the top 65 chr7 peaks (S5 Fig) revealed two longer consensus elements flanking the 5’-TGTG[ta]TGTGTG-3’ motif (Fig 3G, 3H). These consensus regions included several short motifs that are similar to the octanucleotides and often contained AT-rich sequences, such as consecutive sequences of AAA, TTT, or TTTT, as well as an AT mixture. Similar sequence regions existed in the centromeric peak regions of chr13. While these two regions were not identical, they shared nine to ten short motifs while maintaining spacing. In weaker binding sites on chr7 (9–11 reads), these short motifs were present in reduced numbers, indicating that more of the short motifs correspond to stronger binding (Fig 3G, 3H). The centromeres of chromosomes 1, 2, 3, 4, and 5 lacked the long consensus motifs. To verify that these long consensus sequences mediate Ki-67 binding outside the centromeric region, we analyzed a region on chromosome 13 (61,873,135–61,875,156) that exhibited robust, continuous Ki-67 binding (Figure 3I). We found that this region contains fragments of the consensus sequence with the correct spacing (Figure 3J). Therefore, these long consensus motifs function as specific sites for Ki-67 binding to chromosomes.

### GNL2 and MDN1 are mapped to the chromosomal sites where Ki-67 binds

To determine whether these two proteins bind to Ki-67 binding sites on chromosomes, we visualized ChIP-seq data using BAM tracks for each protein. The initial ChIP-seq results (chip1) showed that GNL2 occupied the same binding sites as Ki-67 (Fig 2C–2E). However, the MDN1 occupancy was inconclusive due to its weak read signals. This is not surprising, as MDN1 interacts less closely with Ki-67 than GNL2 does [10]. Subsequently, we amplified the MDN1 reads by exposing twice as many cells to formaldehyde, a cross-linking reagent, for 20% longer. Consequently, MDN1 also aligned with Ki-67 binding regions (chip2). Notably, the number of GNL2 reads decreased drastically in this chip2. Since GNL2 interacts more closely with Ki-67 than MDN1 does [10], the decrease appears to be due to MDN1 or its complex masking GNL2 during the final step of formation.

### GNL2 C-termini assemble the MGM complex

To characterize the MGM complex biochemically, we cloned the GNL2 gene and constructed two plasmids: one that encodes the full-length protein tagged with EGFP and 6xHis tags at the N- and C-termini (Fig 4A, protein G7), and one lacks C-terminal 6xHis tags (protein #1).

**Fig 4.**
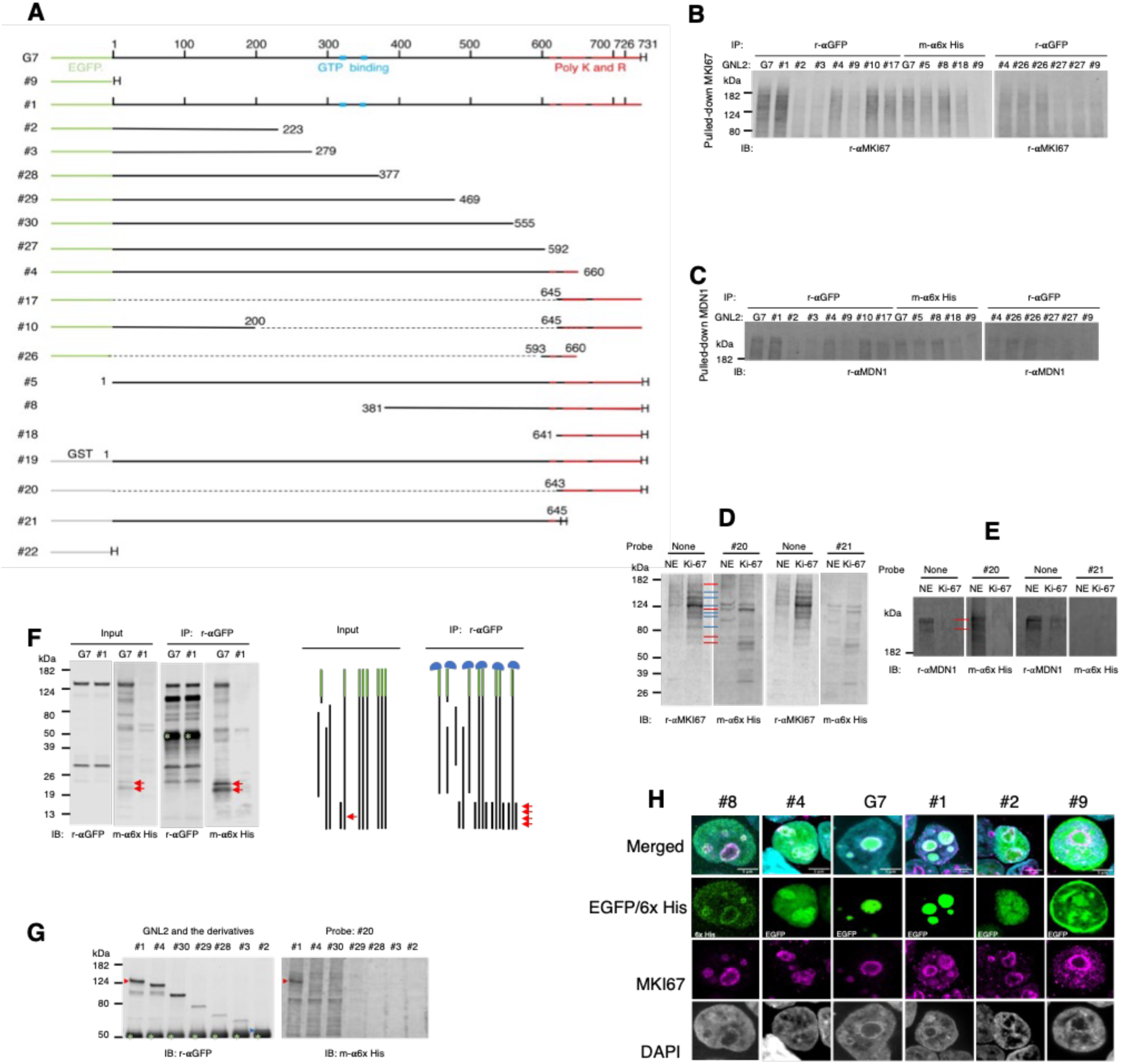
Binding of GNL2 to Ki-67 and MDN1. **A**. Primary protein structures encoded by the cloned GNL2 and its deletion mutants. Black line, GNL2 primary structure; green line, EGFP; gray line, GST; red line, highly charged sequence regions containing lysine and arginine; H, 6xHis tag. **B**. Immunoprecipitation of the endogenous Ki-67 through full-length GNL2 and its deletion mutants. **C**. Immunoprecipitation of endogenous MDN1 through full-length GNL2 and its deletion mutants. **D**. Far Western blots showing a direct binding of the GNL2 C-terminus to Ki-67. Nuclear extracts and purified Ki-67 were subjected to PAGE and transferred to a PVDF membrane. One PVDF membrane was used to detect Ki-67 fragment bands with a rabbit monoclonal antibody. Another membrane underwent slow refolding of the Ki-67 fragments, which were then probed with purified GNL2 C-terminal fragments #20 or #21. Probe binding was visualized using a mouse anti-6x His tag antibody. NE and Ki-67 designate lanes for nuclear extracts and purified Ki-67 fragment lanes, respectively. The red bar shows the Ki-67 binding bands, to which probe #20 bound increasingly in parallel with the Ki-67 fragment levels. The blue bar shows the Ki-67 fragment bands to which #20 did not bind. **E**. Far Western blots showing a direct binding of the GNL2 C-terminus to MDN1. This experiment was performed as described in (D), except that a rabbit polyclonal anti-MDN antibody was used to detect MDN1. The red bar shows the probe #20 binding to MDN1. MDN1 was removed during the Ki-67 purification. **F**. Binding of GNL2 C-termini to full-length GNL2. Two full-length GNL2 proteins (G7 and #1) were transiently expressed in separate HEK293T cultures. The nuclear extracts were used for the Western blot (Input lanes) and for the immunoprecipitation of the tagged GNL2 (IP: r-αGFP lanes) using a rabbit anti-GFP antibody (to the N-terminal tag). The immunoprecipitants were visualized by Western blot using the rabbit anti-GFP and the mouse anti-6xHis antibodies after protein electrophoresis (PAGE) and transfer to a polyvinylidene fluoride (PVDF) membrane. A large quantity of GNL2 C-terminal fragments were precipitated by immunoprecipitation using the N-terminal tag antibody. The green star indicates the rabbit immunoglobulin, and the red arrow indicates the GNL2 C-terminal fragments revealed by the mouse antibody. The right illustration presents an interpretation of the results. The blue cap represents the rabbit anti-GFP antibody, the green bar represents the EGFP tag, and the red arrow represents the GNL2 C-terminal fragments. The GNL2 N-termini were cleaved off during immunoprecipitation. However, they remained interacting with intact GNL2 and co-immunoprecipitated with intact molecules. **G.** The C-terminus-C-terminus interaction between GNL2 molecules. Full-length GNL2 (#1) and its C-terminus-truncated versions were transiently expressed in HEK293T cells and immunopurified using the rabbit anti-GFP antibody. After PAGE of these precipitants, they were transferred to a PVDF membrane. The proteins were slowly refolded, probed with GNL2 C-terminus (#20), and visualized using rabbit anti-GFP and mouse anti-6xHis antibodies. The signals were scanned at 800- and 700-nm windows. The obtained images are shown separately. Green star indicates the immunoglobulin of the rabbit antibody. Protein #2 was present directly above the immunoglobulin, as indicated by a green arrowhead. The red arrowhead shows the probe (#20) binding only to the full-length GNL2. **H**. Nucleolar periphery localization of GNL2 due to interaction between its GNL2 C-terminus and MKI67. GNL2 constructs were transiently expressed in HEK293T cells for two days.

We also created mutant plasmids truncated from the N- or C-termini of the two original plasmids. Then, we transiently expressed the proteins from the plasmids in HEK293T cells and pulled down endogenous Ki-67 using their nuclear extracts and either an anti-EGFP or an anti-6xHis antibody. These antibodies pulled down Ki-67 with nuclear extracts containing the full-length GNL2 but failed to pull down Ki-67 with nuclear extracts containing only the tag (#9) (Fig 4B). Thus, GNL2 binds to MKI67. The fact that the pulled-down Ki-67 was mostly fragmented suggests that Ki-67 is prone to cleavage and that the fragments are more easily immunoprecipitated than the intact protein. Pulldown experiments with truncated constructs revealed that the C-terminus of GNL2 (residues 593–731) binds to Ki-67. This C-terminal binding activity was present in a broad region that could be separated into two regions: residues 593–660 (revealed with protein #4) and residues 645–731 (revealed with protein #10). GNL2^645–731^, which corresponds to IDRs containing blocks of positively and negatively charged amino acids, exhibited stronger binding activity than GNL2^593–660^, which contains fewer charged amino acids. Interestingly, the 645–731 region fragment alone (#17) pulled down less Ki-67 than the same fragment fused to its NLS region (#10). This suggests that full nuclear localization of GNL2 through the NLS is necessary for maximal nucleolar localization. Similarly, endogenous MDN1 was immunoprecipitated (Fig 4C). Next, we examined the interaction between GNL2 and Ki-67 using Far Western blotting. This technology involves two critical steps: refolding of PAGE-separated proteins on a membrane and the specific interaction between a protein and its ligand. The slow refolding process can restore proteins’ enzymatic activity and has been used for many years [75]. Far Western blotting results demonstrated that the C-terminal probe (#20, GNL2^643-731^-6xHis) (see S6 Fig for its purification) bound directly to Ki-67 (Fig 4D). Specifically, the probe increasingly bound to four Ki-67 fragments (highlighted with red lines) as their levels of the Ki-67 fragments increased. Conversely, there were several Ki-67 fragments to which the C-terminal probe did not bind (highlighted with blue lines). These results suggest that the GNL2 C-terminal IDRs bind only to Ki-67 segments with their interacting interfaces. Probe #21 (GNL2^1-645^-6xHis), which had fewer number of positively and negatively charged amino acid residues, bound less strongly to Ki-67 fragments than probe #20. Considering the numerous IDRs present in Ki-67 (Fig 1A), these results imply that the GNL2 C-terminus binds to various Ki-67 peptide regions through IDR-IDR interactions. AlphaFold 3 [76] did not predict an interaction between GNL2 and Ki-67, supporting the conclusion that GNL2 and Ki-67 bind through IDR-IDR interactions. Probe #20 bound to MDN1, but probe #21 did not (Fig 4E). These results suggest that GNL2 binds more strongly to Ki-67 than to MDN1.

Interestingly, we found that the anti-N-terminal EGFP antibody immunoprecipitated C-terminal fragments in addition to the full-length GNL2, from the nuclear extracts in which G7 was transiently expressed from a plasmid. However, we did not obtain the same result with tagless C-terminal GNL2 (#1) because the precipitated C-termini were identified using the anti-C-terminal 6xHis tag antibody (Fig. 4F). Thus, GNL2 forms homocomplex using the C-terminal IDRs. To determine which part of GNL2 the C-terminus binds to, we performed Far Western blotting using purified full-length GNL2 (#1) and its several C-terminus-truncated proteins, as well as probe #20 as the GNL2 C-terminal ligand (Fig. 4G). We found that the probe #20 bound only to the full-length GNL2. Thus, GNL2 forms homomultimers through IDR-IDR interactions at the C-termini. Considering this result alongside the finding that GNL2 binds to Ki-67, we conclude that GNL2 forms homomultimers through IDR-IDR interactions and binds to Ki-67 through the IDR multimers. Similarly, the C-termini of GNL2 may weakly bind to MDN1. However, due to the weak nature of this binding, the alternative possibility of a direct interaction between Ki-67 and MDN1 still exists.

### Ki-67 imports NoLS-containing proteins to the nucleolus

Since the C-terminus of GNL2 overlaps the predicted NoLS, we investigated its role in the nucleolar localization by transiently expressing GNL2 and its mutant proteins (Fig 4H). Observation with a confocal microscope revealed that protein #8 (GNL2^381-731^) precisely colocalized with Ki-67 at the nucleolar periphery. Protein #4 (GNL2^1-660^) colocalized with Ki-67 but was also present separately from Ki-67 within the nucleolus while remaining in the nucleus at high levels. The two full-length proteins (G7 and #1) colocalized with Ki-67 and were also present separately from Ki-67 within the nucleolus but, unlike #4, did not remain in the nucleus. In contrast, protein #2 (GNL2^1-223^), which has an NLS but lacks the C-terminus and most of the GNL2 portion, accumulated in the nucleus and reached the nucleolus. However, it did not concentrate at the nucleolar periphery. Tag #9 (EGFP-6xHis) also entered the nucleolus through the nucleus without concentrating and was predominantly present in the cytoplasm due to the absence of an NLS. These results demonstrate that binding activity of the C-terminus of GNL2 (GNL2^593-731^) to Ki-67, primarily via the predicted NoLS, is essential for GNL2 localization in the nucleolus. This can be generalized to the nucleolar localization of other proteins with an NoLS.

### The MGM complex is involved in the export of pre-60S particles through the nucleolar periphery

Next, we knocked down these three genes to understand the role of the MGM complex at the nucleolar periphery. Knocking down MKI67 decreased the levels of GNL2 and MDN1 levels in the nucleolus and the surrounding area while increasing their cytoplasmic levels (Fig 5A11, 5A15, 5A14, 5B, 5C).

**Fig 5.**
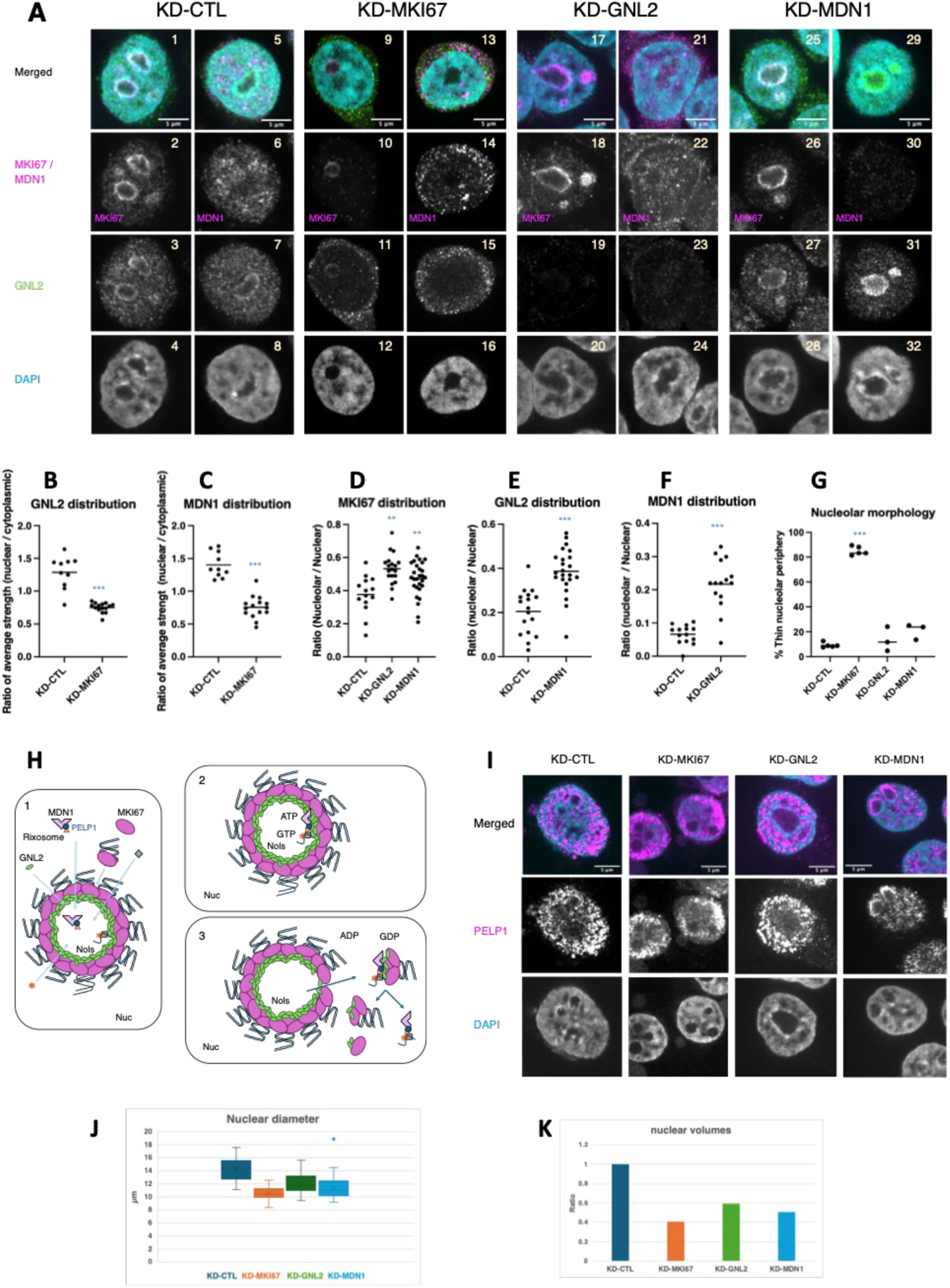
Morphological changes caused by knockdown of MKI67, GNL2, and MDN1. **A**. Images resulting from the knockdowns. Only the merged images are displayed in color. **B–G**. Results of the statistical analysis of the images in (**A**). T-test: **, P < 0.001; *** P < 0.0001. **H**. A model of the import of nucleolar proteins, protein complex, and nucleoproteins and the export of the pre-60S particles through the nucleolar periphery. **H1**. Synthesized pre-rRNA and imported proteins, protein complexes, and nucleoproteins form particles with GNL2 present on the nucleolar peripheral Ki-67. **H2**. The GNL2 particles then recruit the rixosome complex (the RIX1 complex plus MDN1) to the nucleolar periphery. **H3**. The recruited MDN1 then disintegrates the peripheral MKI67 complex and releases/exports the pre-60S particles into the nucleus. **I**. Confirmation of the role of MDN1 in the export of pre-60S particles. Knocking down *GNL2* resulted in the accumulation of PELP1 within the nucleolus. However, knocking down *MDN1* immobilized PELP1 on the nucleolar periphery. PELP1 serves as a marker for pre-60S particles. **J**. Length of the longer nuclear diameter. Forty nuclei from two independent cultures were measured for each knockdown sample. T-test: all three were < 1E-7 compared to the control. The diameter of MKI67-knockdown nuclei was smaller than those of GNL2- and MDN1-knockdown nuclei, respectively. T-Test: P < 1E-4 in CTL vs. KD-MKI67, and P < 0.05 in CTL vs. KD-GNL2 or KD-MDN1. **K**. The nuclear volume ratios of MKI67-, GNL2-, and MDN1-knockdown nuclei to control knockdown nuclei were estimated using the following formula: ((4π(L_T_/2)³/3)/((4π(L_C_/2)³/3) = (L_T_/L_C_)³, where L_T_ and L_C_ are the average longer diameters of the knockdown and control nuclei, respectively.

This phenotype may result from interrupting the nucleolar import of GNL2 and MDN1, increasing the export of pre-60S particles, or both. Conversely, knocking down *GNL2* increased nucleolar size, as well as Ki-67 and MDN1 levels, in the nucleolus (Fig 5A17, A18, A21, A22, D, F). Similarly, knocking down *MDN1* increased nucleolar size and both Ki-67 and GNL2 levels in the nucleolus (Fig 5A25, A26, A29, A31, D, E). The accumulation of Ki-67 suggests that both GNL2 and MDN1 are necessary for releasing Ki-67 from the nucleolus. The accumulation of MDN1 by *GNL2* knockdown and the accumulation of GNL2 by *MDN1* knockdown suggest that both proteins are necessary for exporting pre-60S particles from the nucleolus. These results are summarized in Fig. 5H, which includes the most likely scenario. Briefly, imported proteins and RNAs, as well as synthesized and matured rRNAs form a complex with GNL2 on Ki-67. This complex corresponds to the *S. cerevisiae* Nog2 particle, i.e., GNL2 particles. Next, pre-60S particle forms with the rixosome joining. MDN1, which is present on the pre-60S particle, then disintegrates the Ki-67 layer. This allows the Ki-67 and the particle to exit the nucleolus. To support this model, we investigated the behavior of PELP1, which is another pre-60S particle subunit that binds directly to MDN1 [41, 42, 46]. Knocking down *GNL2* increased PELP1 levels within the nucleolus (Fig 5I), as was observed with MDN1 (Fig 5A21, 5A22). Importantly, knocking down MDN1 trapped PELP1 and increased its levels on the nucleolar periphery. Thus, we conclude that MDN1 is directly involved in exporting pre-60S particles to the nucleus, and that GNL2 is involved in forming particles that include MDN1 at the periphery.

Additionally, knocking down MKI67 produced structural phenotypes. It reduced nucleolar peripheral DNA and formed clumps of less-dense DNA of various sizes that were dispersed throughout the nucleus (Fig 5A12, 5A16, 5G). These results suggest that Ki-67 plays a significant role in tethering chromatin to the nucleolar periphery, where silenced genes tend to localize [77–80]. Furthermore, the knockdown decreased the nuclear size of cells by an average of 26% in terms of the longer diameter. This shrinkage enabled us to estimate a maximum 60% decrease in volume (Fig 5J, 5K; S7 Fig). Since small nucleoli caused by MKI67 knockdown are associated with decreased pre-RNA levels and cell size in HeLa cells [8], we can infer that protein synthesis is reduced in MKI67-knockdowned HEK293T cells. Knocking down GNL2 or MDN1 also decreased their nuclear size. However, the decrease in these cells exhibited greater deviation than the decrease in Ki-67 knockdown cells.

### Depleting Ki-67, GNL2, and MDN1 decreases the levels of transcripts involved in ribosome biogenesis; depleting Ki-67 increases the levels of numerous neuron-specific transcripts

We examined the efficacy of knocking down the *MKI67*, *GNL2*, and *MDN1* genes. Three days after transfecting siRNA for each gene, the transcript levels of the *MKI67*, *GNL2*, and *MDN1* gene decreased by 70–80% without secondary effects (Fig 6A).

**Fig 6.**
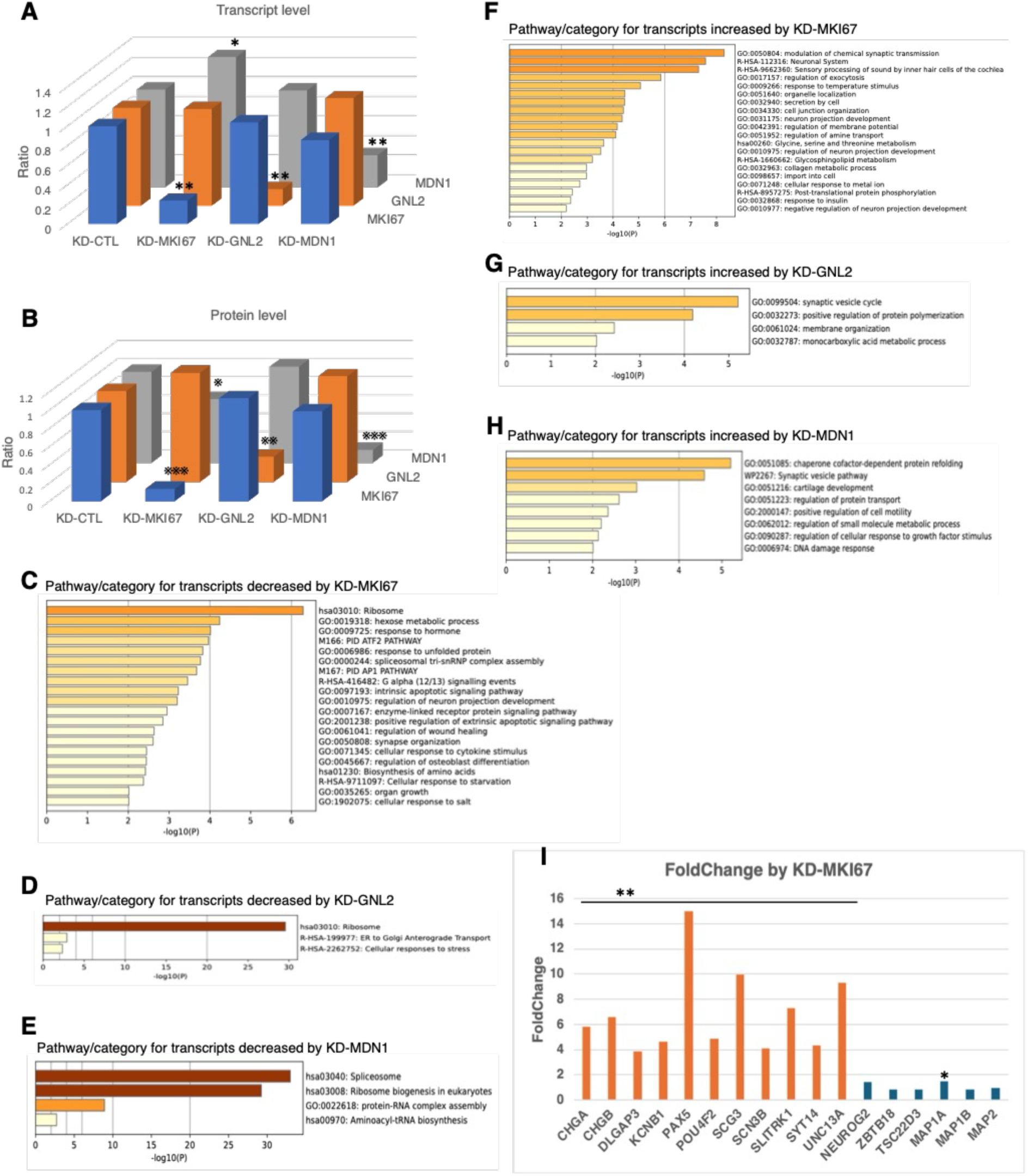
Efficacy of *MKI67, GNL2*, and *MDN1* knockdown and its effect on genomic transcripts. **A**. Levels of the *MKI67, GNL2*, and *MDN1* transcripts in the knockdown cultures. These data result from the RNA-seq experiments. Each transcript level is presented as a ratio relative to that of the control culture (KD-CTL). Padj: *, <0.001; **, < 1E-37. **B**. Protein levels in the knockdown cultures. Protein levels were determined by Western blotting in four independent cultures. P-value by T-test: *, p < 0.05; **, p < 0.01; and ***, p < 1E-05. **C–E**. Metascape bar graphs showing genomic transcripts downregulated by *MKI67, GNL2*, and *MDN1* knockdown. **F–H**. Metascape bar graphs showing genomic transcripts upregulated by *MKI67, GNL2*, and *MDN1* knockdown. **I**. Examples of expression levels in neural cell-specific transcripts that were either upregulated or unaffected by *MKI67* knockdown. The original data for **C**–**I** are shown in S5–S7 Tables.

The cognate protein level decreased by the same extent or more without affecting the levels of the other two non-targeted proteins (Fig 6B). These results demonstrate that knocking down the *MKI67*, *GNL2*, or *MDN1* gene depletes only the targeted protein, thereby inducing changes in its downstream events. However, knocking down MKI67 slightly decreased the level of the MDN1 protein. This suggests the possibility that knocking down MKI67 may cause downstream changes via the reduced MDN1 level. Given these findings, we next investigated the impact of each protein depletion on genomic transcripts using RNA-seq. The results revealed that each depletion up- and down-regulated numerous transcripts (S8 Fig). Subsequent analysis of the data using Metascape [81] with a fold change cutoff of ≥ 2 revealed that these depletions significantly decreased transcript levels in the ribosome and ribosome synthesis categories (Fig 6C–6E; S5–S7 Table). The strongest knockdown effect was observed with MDN1 depletion, including *rRNA*s, *SNORA*s, *SNORD*s, and many spliceosome transcripts such as *RUN1* (Fig. 6E; S9 Fig). Furthermore, MDN1 depletion most significantly increased the levels of the heat shock protein transcripts *HSPA1A* and *HSPA1B* (S5–S7 Table). These results suggest that MDN1 depletion most strongly arrests the pre-60S particle processing, causing pre-60S particle collapse. In contrast, increased transcripts were less common among the three protein depletions (Fig 6F–6H; S5–S7 Table). However, Ki-67 depletion exhibited the strongest effect, increasing the levels of 100 different transcripts. The top 20 transcripts included 17 neuron-specific transcripts, including those involved in the modulation of chemical synaptic transmission, the neuronal system, and the sensory processing of sound by the inner hair cells of the cochlea. These transcripts were *PAX5, SCG3, UNC13A, SLITRK1, CHGB, CCDC92B (*AC005696.4), *CHGA, POU4F2, KCNB1, SYT14, SCN3B, DLGAP3, AP3B2, LRATD1 (FAM84A), SYN1, VGF, and CPXM1 (CPX1)*. These transcripts increased 15- to 3-fold compared to control levels (Fig 6I; S5 Table), suggesting that Ki-67 plays a significant role in silencing the expression of neuron-specific transcripts. Nevertheless, Ki-67 depletion did not increase *NEUROG2*, *ZBTB18*, *TSC22D3*, *MAP1B*, *ISL2*, and *MAP2* levels. Furthermore, *NEUROG2* and *MAP2* levels were significantly low regardless of Ki-67 depletion. These two genes are important members of the cascade gene network necessary for the neural differentiation of neural progenitor cells [71, 82]. and low *NEUROG2* and *MAP2* transcript levels are consistent with the absence of neuronal structures in HEK293T cells. These results reveal two systems that control gene expression involved in neural differentiation and neuronal activity: the Ki-67-dependent system and the cascade-dependent system. Interestingly, the top 20 transcript group included the *NEK7* transcript. While not neuron-specific, this transcript encodes a kinase necessary for microtubule nucleation at the centrosomes, as well as for cell cycle progression at prophase and neuronal morphogenesis [83, 84].

### Ki-67 reduces transcript levels by either degrading transcripts or silencing gene expression

The neuron-specific transcripts that increased through Ki-67 depletion were transcribed from the genes located in the black, gray, or white zones of the GRCh38/hg38 genomic map. This suggests that Ki-67 is involved in gene regulation through multiple mechanisms. Furthermore, Ki-67 colocalized with YTHDC2 and CCR4-NOT in areas near the nucleolus (5) or nuclear membrane (Fig 7L), indicating the possibility that Ki-67 may be involved in transcript degradation.

**Fig 7.**
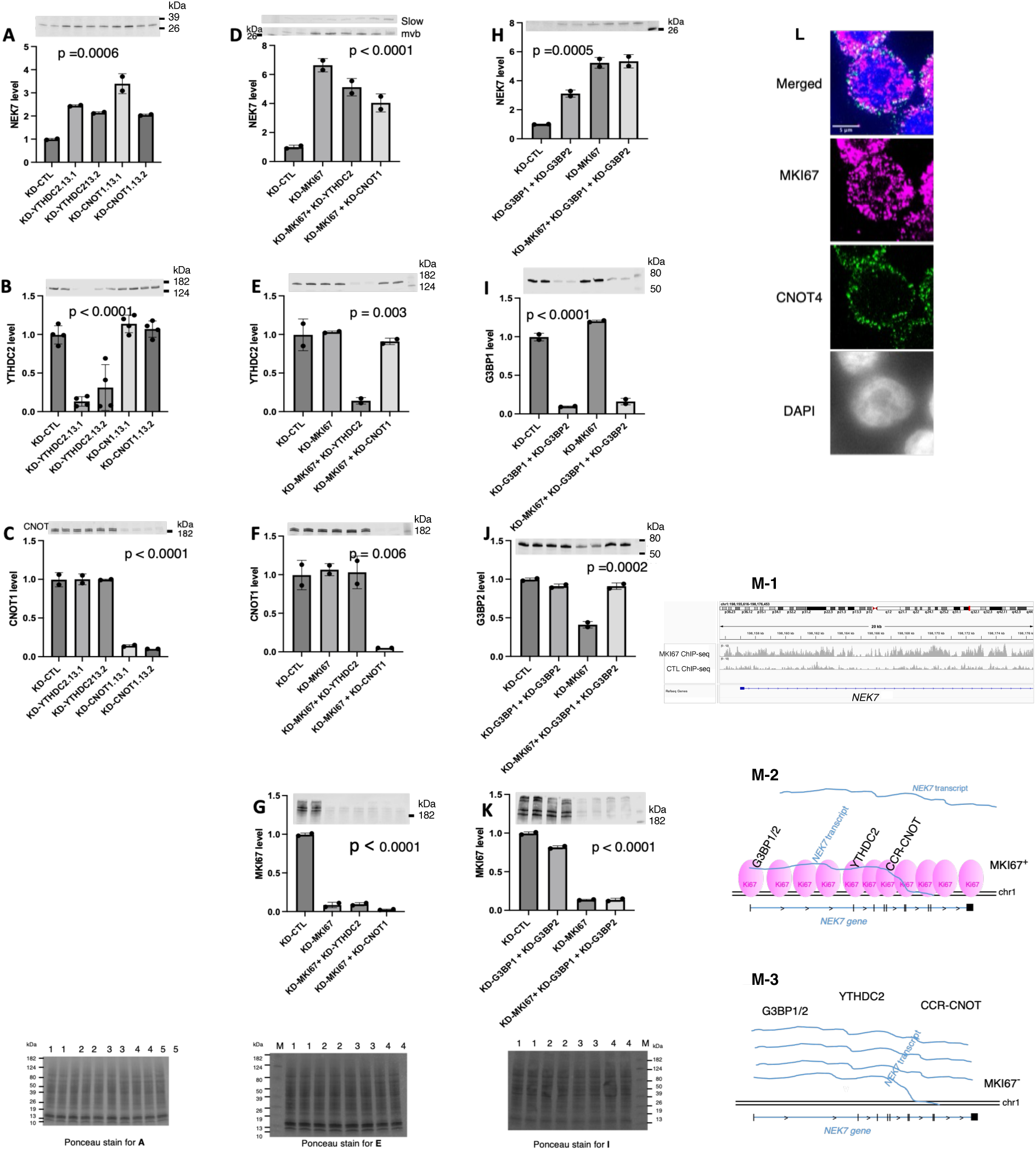
Increase in NEK7 protein levels through the knockdown of *YTHDC2, CNOT1, G3BP1/2*, and *MKI67*. *CNOT1* encodes a scaffold protein of the CCR4-NOT system. A one-way analysis of variance (ANOVA) test was conducted to determine the statistical significance of the results, and the p-values are shown in each figure panel. **A**. Increase in NEK7 protein levels through knockdown of either *YTHDC*2 or *CNOT1*. **B–C**. Confirmation of the genes’ knockdown. **D**. Increase in NEK7 protein levels through *YTHDC2* or *CNOT1* knockdown, along with *MKI67* knockdown. **E–G**. Confirmation of gene knockdown for (D). **H**. Increase in NEK7 protein levels through *G3BP1* and *G3BP2* knockdown, as well as *G3BP1* and *G3BP2* knockdown along with *MKI67* knockdown. **I–K**. Confirmation of gene knockdown for (H). For all Western blots in these experiments, 50 µg of extract protein was loaded into one well of a PAGE gel. The Ponceau staining image for each experiment is presented in the lane below. **L**. Colocalization of Ki-67 with CNOT4, a subunit of the CCR4-NOT system, at the nuclear periphery. Images of Ki-67 and CNOT4 were obtained by adjusting the Z position from the center of the nucleolus to a position where CNOT4 could be clearly observed. **M-1**. Binding of Ki-67 to the 5’ region of the NEK7 gene. **M-2.** A model illustrating the degradation process of the NEK7 transcripts (blue lines). Ki-67 (pink) collects NEK7 nascent transcripts and transfers them to the G3BP1/2-containing stress granules, YTHDC2, and the CCR4-NOT system, This results in NEK7 transcripts degradation. **M-3**. However, the absence of Ki-67 results in NEK7 protein hyperproduction.

Consequently, we knocked down the *YTHDC2*, *CNOT1*, and *G3BP1* genes and investigated the effect on *NEK7*, a transcript regulated by Ki-67 depletion. As expected, knocking down each gene increased NEK7 protein levels two- to three-fold (Fig 7A–7C, see S10 Fig for full blots). However, the increase never exceeded the level caused by Ki-67 depletion when Ki-67 depletion was conducted alongside the knockdown of these genes (Fig 7D–7K). These results suggest that Ki-67 is necessary for degrading mRNA through YTHDC2 and the CCR4-NOT complex and for storing mRNA through G3BP1-containing stress granules. Therefore, Ki-67 likely captures *NEK7* transcripts as soon as they are transcribed and delivers them to YTHDC2, the CCR4-NOT complex, and G3BP1-containing stress granules (Fig 7M-1). This mechanism prevents *NEK7* transcripts from accumulating excessively (Fig 7M-2). Conversely, Ki-67 depletion results in the loss of the delivery, leading to high *NEK7* transcript and protein levels (Fig. 7M-3).

Unlike NEK7, depletion of YTHDC2, CNOT1, G3BP1, or FMR1 did not increase UNC13A protein levels (Fig 8A–E; see S11 Fig for full blots).

**Fig 8.**
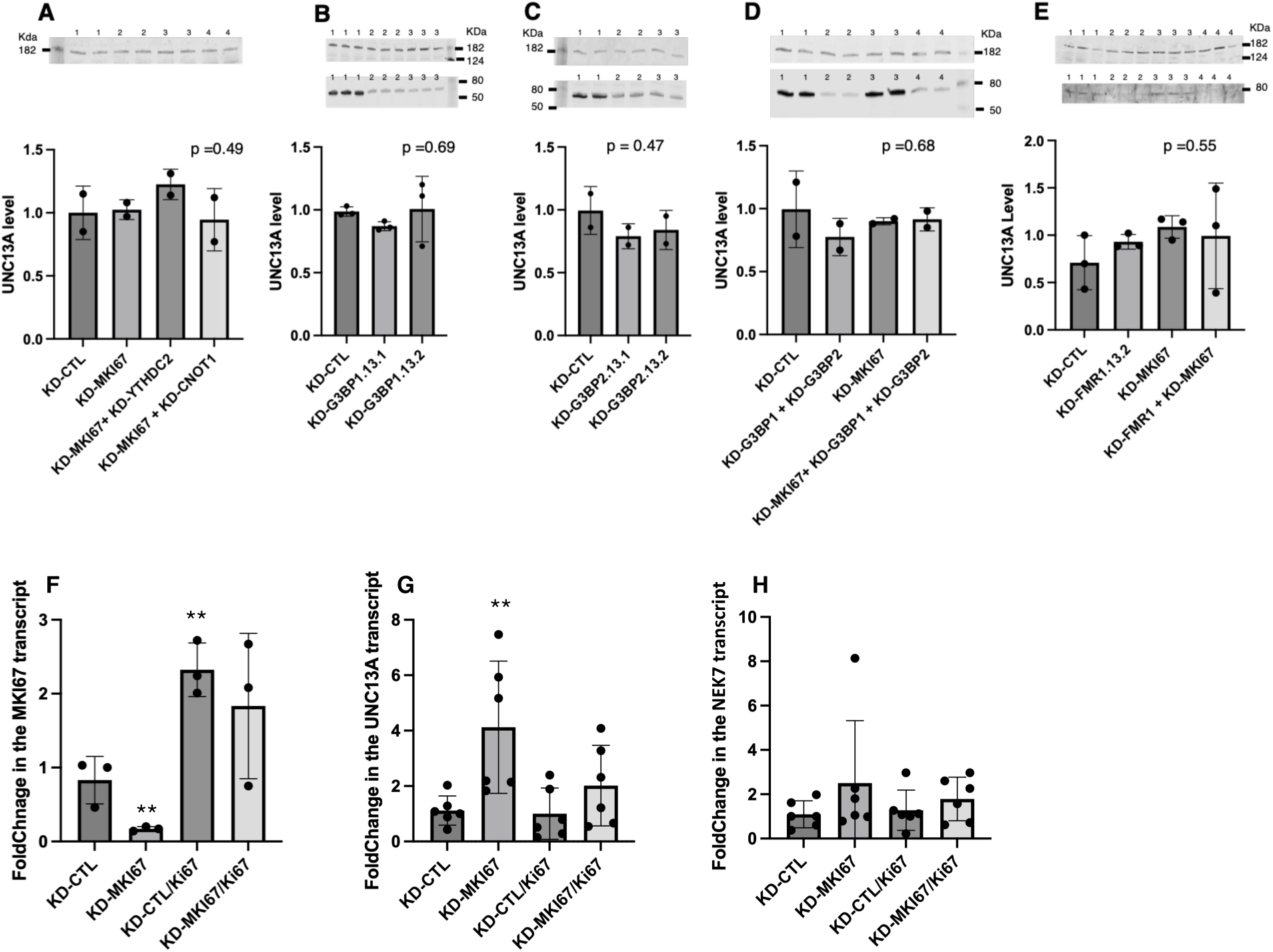
Evidence for the local translation of *UNC13A* transcripts. **A–E**. Knockdown of *YTHDC2, CNOT1, G3BP1/*2, or *FMR1* did not increase UNC13A protein levels. **A, D, E**. Similarly, knockdown of *MKI67* did not increase UNC13A protein levels. The upper Western blots show the UNC13A protein levels, and the lower Western blots confirm the depletion of the targeted proteins. A one-way analysis of variance (ANOVA) test was conducted to determine the statistical significance of the results. **F–G**. Presentation of qPCR results in experiments involving MKI67 knockdown and Ki67 overexpression. The cultures were stably transfected with a plasmid encoding EGFP or EGFP-Ki67. KD-CTL, KD-MKI67, KD-CTL/Ki67, and KD-MKI67/Ki67, respectively, indicate cultures knocked down with control siRNA, siMKI67 RNA (expressing EGFP), control siRNA (expressing EGFP-Ki67), and siMKI67 RNA (expressing EGFP-Ki67). Each dot represents an independent culture. **F**. Confirmation that Ki-67 depletion due to MKI67 knockdown and increased *MKI67* transcript levels due to EGFP-Ki67 plasmid transfection. **G**. Confirmation that Ki-67 depletion increases UNC13A transcript levels, while increased MKI67 decreases them. **H**. Changes in NEK7 transcript levels due to Ki-67 depletion and hyper-expressed MKI67. T-test: **, p < 0.05 compared to control cultures (KD-CTL).

Additionally, contrary to the RNA-seq results, Ki-67 depletion did not increase *UNC13A* transcript levels. These results call into question whether Ki-67 depletion truly increases *UNC13A* transcript levels. Consequently, we performed qPCR analysis, which confirmed that Ki-67 depletion increased *UNC13A* transcript levels (Fig 8F, 8G; S8 Table, S9 Table) and demonstrated that overexpressing Ki-67 (in the form of EGFP-Ki67) decreases the elevated *UNC13A* transcript level back to nearly the control level. These results provide the proof that Ki-67 is responsible for silencing *UNC13A* gene expression and that its depletion enables the gene expression. Therefore, both observations stand: Ki-67 depletion causes high UNC13A transcript levels yet does not increase its protein levels. This discrepancy between transcript and protein levels is attributed to a unique feature of the HEK293T cell line—it does not form the dendrites and axons required for *UNC13A* translation. We also analyzed *NEK7* transcript levels using qPCR and found that they tended to be antiparallel to *MKI67* transcript levels, as anticipated (Fig 8H; S8 Table, S9 Table). In summary, Ki-67 functions at two different steps to decrease protein levels. First, it promotes mRNA decay, as observed with *NEK7*. Second, it silences gene expression, as observed with *UNC13A*.

### Transcripts upregulated by Ki-67 depletion include those that promote neuronal development

Ki-67 depletion increases the levels of numerous neuron-specific transcripts, including *NEK7, POU4F2,* and *PAX5* that increased four- to 15-fold (S5 Table). *NEK7* is necessary for prometaphase progression [83] and dendrite growth, branching, spine formation, and morphogenesis [84]. *NEK7* upregulation also promotes cell proliferation in gastric cancer cells [85]. *POU4F2* is necessary for forming cells that resemble retinal ganglion cells (RGCs) [86]. *PAX5* is necessary for brain development, axonal guidance, neurotransmitter regulation, and neuronal adhesion [87, 88]. Thus, high levels of *NEK7, POU4F2*, and *PAX5* transcripts due to Ki-67 depletion enable cells to proliferate and develop into neurons. To gain deeper insight into the proliferation mechanism in Ki-67-depleted HEK293T cells, we searched the RNA-seq dataset of these cells for additional candidate transcripts that contribute to neuronal development. We identified several apoptosis-related transcripts that were increased by Ki-67 depletion, including BNIP3 and DDIT3; however, qPCR analysis of these increases was inconclusive (S10 Table) [89–92]. In contrast, *SLC7A2* levels increased by Ki-67 depletion, and qPCR analysis approved the increase and showed that Ki-67 overexpression (EGFP-Ki67) reduced the elevated levels to control levels (S10, S11 Table).

*SLC7A* encodes a transporter of cationic amino acids (e.g., L-arginine, L-lysine, and L-ornithine), which activate mTORC1. Therefore, high levels of *SLC7A2* caused by Ki-67 depletion can promote protein synthesis and proliferation. This effect has been observed in several studies, though it occurs independently of the Ki-67 effect [93–96]. Taken together, these results suggest that high levels of *NEK7*, *POU4F2*, *PAX5*, and *SLC7A2* caused by a lack of Ki-67 drive protein synthesis to levels similar to those in proliferating cells. This enables neural progenitor cells (NPCs) to differentiate and develop into neurons. Sufficient Ki-67 levels promote nucleolar protein import and pre-60S particle export. This results in NPC self-renewal and silencing of neuron-specific gene expression across the genome (Fig 9D). However, when Ki-67 is lost, the levels of these four transcripts increase in NPCs, leading to the maintenance of protein synthesis, followed by differentiation and development. Neurodevelopmental events in humans are initiated during embryogenesis and are mostly completed by birth, though NPCs persist to some extent. Therefore, shortly after birth, most neurons in the brain are post-mitotic [97]. These neurons then migrate to their proper locations and form functional circuits [98]. Thus, the absence of Ki-67 during the differentiation and post-mitotic development is crucial for completing and maintaining neuronal circuits.

**Fig 9.**
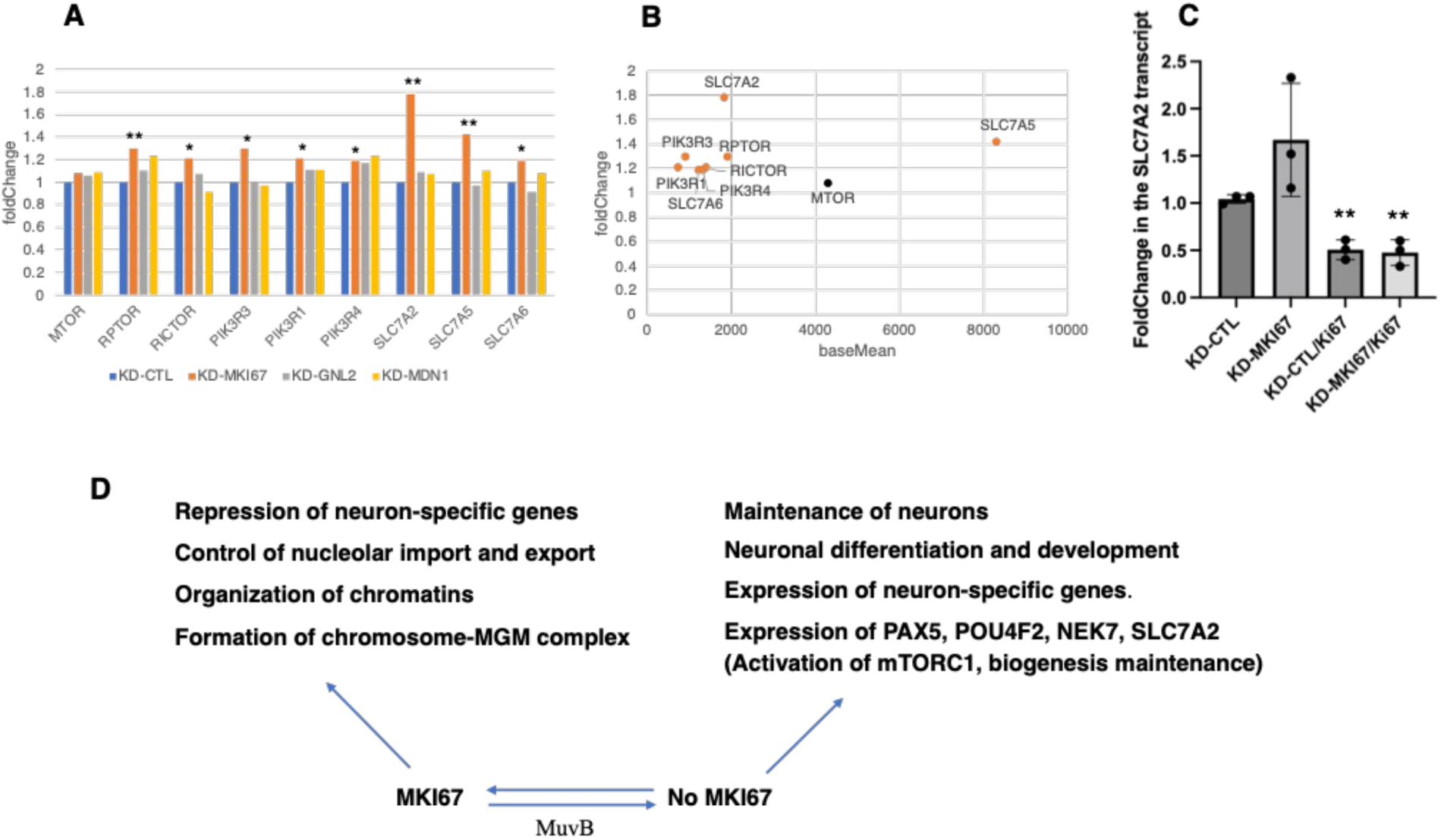
The presence or absence of Ki-67 activates various cell cellular processes. **A**, **B**. mTORC1-activating transcripts whose levels increased by MKI67 depletion. The data in these panels are derived from the RNA-seq dataset. Padj value: *, < 0.05; **, < 0.001. **C**. Ki-67 dependent changes in *SLC7A2* transcript levels examined by qPCR. This experiment was performed as described in Fig 8 F–G. T-test: **, p < 0.005. **D**. The proposed MKI67 functions in NPCs, as well as during differentiation and in neurons. See the text for more information.

### Depletion of MDN1 results in low levels of respiratory complex transcripts

We observed an acidification in MDN1- and Ki-67-depleted cultures but not in GNL2-depleted cultures or the control. Acidification began two days after the depletion and persisted for the next three days (Fig 10A; S12 Fig).

**Fig 10.**
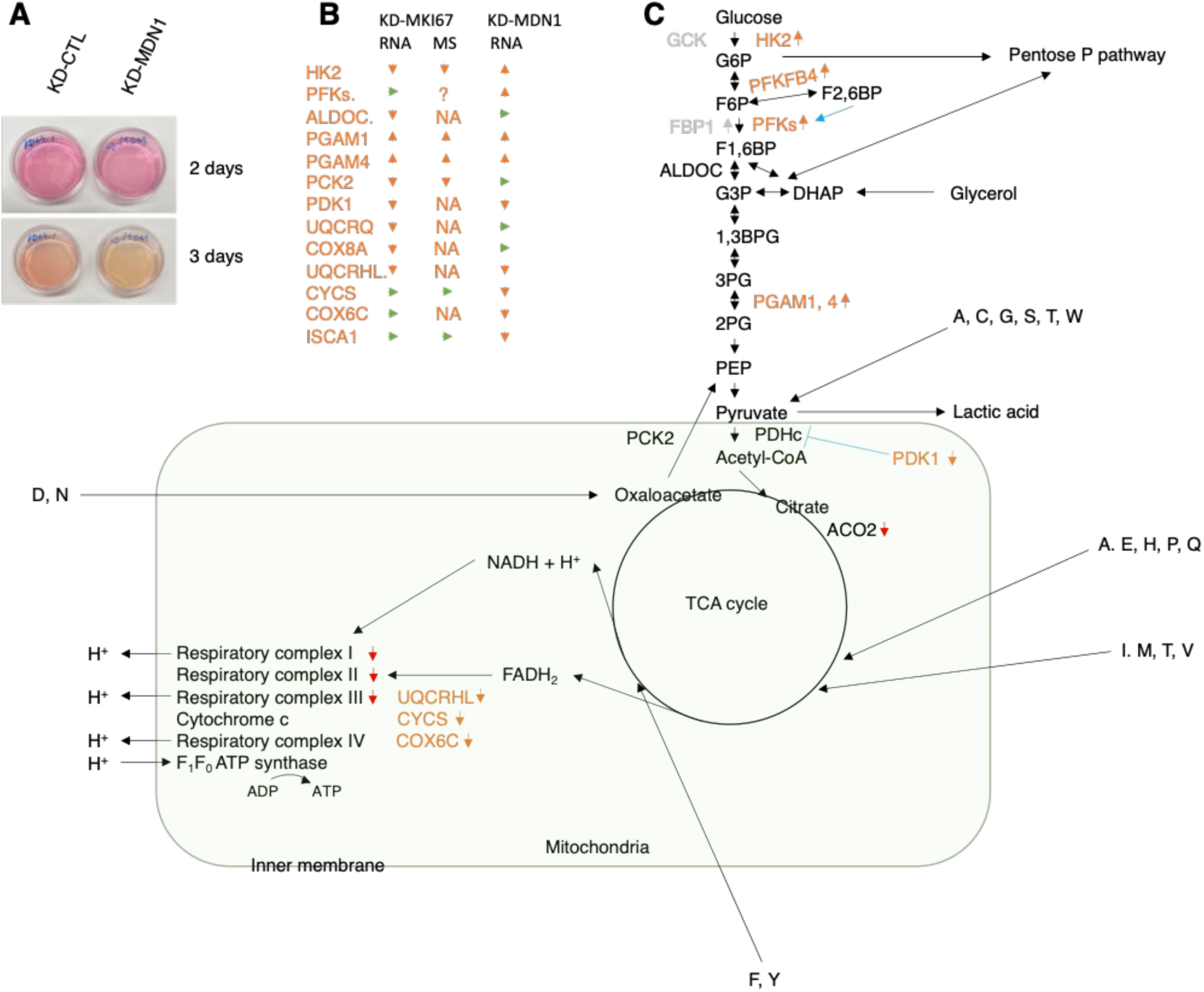
The shifts in transcript levels involved in respiration and glycolysis caused by depleting MDN1 or Ki-67. A. pH decrease in an MDN1-knocked down HEK293T culture. **A**. HEK293T cells (4x10⁵) in a 2.8 ml volume were knocked down for MDN1 and the control in a 33-mm Matsunami dish. The cultures were then incubated for three days. **B**. Arrowheads indicate changes in expression levels obtained by RNA-seq and MS experiments three days after MKI67 or MDN1 knockdown. Orange arrowhead indicates an increase or a decrease, while green arrowhead indicates no change. The “NA” indicates that the data is unavailable. MDN1 depletion decreased the number of respiratory enzyme transcripts more than Ki-67 depletion did. Depletion also decreased the transcript levels of a few glycolytic enzymes, while Ki-67 depletion weakly increased the transcript levels of a few glycolytic enzymes. **C**. Predicted alterations in the catalytic activities in respiration, the TCA cycle, and glycolysis in MDN1-depleted cells. These alterations are also based on transcript alterations obtained from the KD-MDN1 RNA-seq dataset. Amino acids are denoted by one-letter abbreviations. Orange arrowhead was used as in (B). Black arrow is used to denote reaction fluxes, and blue line indicates enzyme activation or inhibition. Red arrowhead shown next to the respiratory complexes and aconitase 2 (ACO2) signifies a predicted decrease in the catalytic activity due to the downregulation of ISCA1 transcripts.

Acidification was more intense in the MDN1-depleted cultures, which was most evident on the final day of the experiment. Searching the RNA-seq dataset revealed that many transcripts were involved in this phenomenon. MDN1 depletion severely decreased the levels of the mitochondrial transcripts *MT-RNR1* and *MT-RNR2* (S9A Fig), as well as the levels of many transcripts encoding respiratory proteins, including *ISCA1* and *CYCS* (Fig 10B, 10C; S12 Table). ISCA1 integrates iron-sulfur centers, which are essential for electron transfer, into respiratory complexes I, II, and III, as well as the TCA cycle enzyme aconitase 2 [99]. Thus, despite the twofold decrease, the decrease in *ISCA1* was predicted to severely impair respiration by causing simultaneous impairments in these four metabolic steps (Fig 10C). Meanwhile, the levels of transcripts encoding glycolytic enzymes (*HK1, HK2*, *PFKP, PFKL, PFKM*, *PGAM1, and PGAM4*) increased slightly, contributing to increased lactic acid production (S12 Table). These results suggest that MDN1 plays a key role in preventing excess energy production when ribosome biogenesis, a highly energy-consuming process, is low. This MDN1-mediated regulatory mechanism acts in the opposite direction of the energy-dependent nucleolar silencing complex (eNoSC). The eNoSC reduces ribosome biogenesis by repressing rRNA transcription when the energy sources are scarce (e.g., when glucose levels are low) [100]. Unlike with MDN1 depletion, Ki-67 depletion decreased *HK2* levels and increased *PGAM1* and *PGAM4* levels (S12 Table). This suggests that, without Ki-67, cells maintain the flow of sugar carbon skeletons to the pentose phosphate shunt in order to synthesize nucleic acids for RNA and protein production.

### MDN1 binds almost exclusively to genes transcribed by Pol III

MDN1 binds to promoter regions independently of Ki-67 and GNL2. Specifically, MDN1 bound to the 5’ ends of protein-coding genes, such as *DAB2IP*, *PAK4*, *RFX1*, and *SGSM1* (S13A, 13B Fig), as well as to the 5’ ends of Pol III-dependent genes, such as the *RNY1/3/4*, *RNU6-1/2/7/8/9, RMRP*, and *RN7SK* gene (S13C–F Fig). However, MDN1 bound to the latter group over ten times more than the former. On these RNA genes, MDN1 also bound weakly to the 3’ ends of the *RNY3* and *RNU6-2* genes, as well as to the entire *RMRP* and *RN7SK* genes. Nevertheless, depleting MDN1 did not significantly alter the transcript levels of these genes more than transcript levels of RNA genes to which MDN1 did not bind (S14A, 14B Fig). These results suggest that MDN1 activates these Pol III-dependent genes or plays a role other than that of a transcription activator, e.g., a transcript stabilizer. Another possible role of MDN1 is selecting Pol III over Pol II to ensure precise termination of transcription at the 3’ end of these small genes [47, 101–103]. Interestingly, although several *RNU* genes exist, MDN1 binds only to *RNU6* genes, which encode transcripts that recruit Mg²⁺ ions and splice pre-mRNA [47]. This feature may indicate MDN1’s role in binding to Pol III-dependent genes.

## Discussion

### The significance of Ki-67 binding to entire chromosomes

This report provides evidence that Ki-67 binds to chromosomes genome-wide through its nucleotide specificity. MacCallum and Hall reported on Ki-67’s affinity for AT-rich nucleotide sequences [13]. While we agree with their findings, we found that AT-rich sequences alone are not sufficient for binding and identified the specific nucleotide sequences that enable Ki-67 to bind to chromosomes. van Schaik et al. Ki-67’s genome-wide binding using the pA-DamID mapping method [18]. This method identifies Ki-67 binding sites by determining the GATC sites that the protein A-Dam enzyme fusion protein methylates in loci adjacent to the anti-Ki-67 antibody. This method has advantages over classical Ki-67 ChIP-seq: it is simpler and more efficient at tracking changes in Ki-67 binding sites throughout the cell cycle. However, their results regarding Ki-67 binding regions differed greatly from ours. For example, pA-DamID-identified Ki-67 binding sites on chr2 in randomly growing cultures were narrower than ours. Their underestimation of the binding regions appears to result from the features of their method itself, pA-DamID. There may be three reasons for their underestimation. First, since pA-DamID indirectly identifies Ki-67 binding sites by methylating GATC nucleotide sequences adjacent to these sites, pA-Dam may not methylate regions of the chromosome that are densely and thickly coated with Ki-67. Second, the pA-Dam cannot detect GATC sites where N6-adenosine has been methylated by human methylases [104]. Third, the anti-Ki-67 antibodies used by van Schaik’s group and our group had different binding features for Ki-67. Complete coating during interphase is important, as it is during mitosis [6]. This importance during interphase was suggested by chromosome clumping caused by Ki-67 depletion (Fig 5A12,5A16). Other groups recognized the importance of nearly complete coating via observation of DNA damage [8, 31], although this phenotype does not appear until prophase. The appearance of this phenotype coincides with the removal of cohesins, which bundle the duplicated double-helical strands during interphase and begin to be removed from the duplicated strands during prophase [105, 106]. Adding the DNA strand breaker phleomycin, which increases the frequency of this damage, more clearly demonstrates that the damage occurs only during mitosis, from prophase through anaphase [31]. Cohesins are necessary for repairing DNA double-strand breaks (DSBs) [107]. This suggests that clumping with bare chromosomes due to Ki-67 depletion may increase the probability of intra- and interchromosomal entanglements during interphase, which can subsequently cause DSBs during mitosis.

### The roles of Ki-67 in importing nucleolar proteins and processing pre-60S particles on the nucleolar periphery

We found that Ki-67 serves as a receptor of GNL2 nucleolar import, probably also of over 4,500 nucleolar proteins [108]. Of these proteins, those containing NoLS motifs should actively enter the nucleolus through the binding to peripheral Ki-67 and then migrate to their destinations. In the cases of proteins lacking an NoLS, they can actively enter the nucleolus by using NoLSs available from subunits of their complexes. For example, our analysis using the Nucleolar Localization Sequence Detector revealed that the rixosome complex contains two proteins without NoLSs (PELP1 and WDR18) and four proteins with NoLSs (MDN1, NOL9, SENP3, and TEX10) [33, 34]. After imported, the proteins would be recruited to the destinations through stronger affinity for special proteins, such as POLI and nucleophosmin 1(NPM1), or DNA/RNA within the nucleolus [109]. Additionally, we found that Ki-67 maintains pre-60S particles in the nucleolus and, subsequently allows them to exit the compartment through Ki-67 via a GNL2-and MDN1-involved mechanism. Therefore, both nucleolar import and export are important processes only in human and other species’ cells that contain Ki-67 [40]. However, Ki-67-depleted HEK293T cells proliferate against the expectation deriving from the function of Ki-67 at the nucleolar periphery. The reason why the Ki-67-depleted cells can proliferate is likely due to less presence of obstacles, i.e., Ki-67 and chromatins, for free diffusion of nucleolar proteins in and pre-60S particles out of the nucleolus. The smaller nucleolar size should also help these free diffusions, while it also indicates that Ki-67-depleted cells produce less pre-rRNA and lower protein synthesis [8], implying that they proliferate slowly than control cells. The biogenesis of ribosomes in Ki-67-less nucleoli is important because the nucleoli lacking Ki-67 but still must function for ribosome biogenesis during developing neurons, neurite morphogenesis, and the long-term maintenance of mature neurons [110].

### The disappearance of Ki-67 plays a key role in promoting neuronal differentiation, development, and maintenance

Chromatin interacts with the nucleolus through NADs. The membrane-less structure of the nucleoli complicates NAD investigations. However, new technologies are revealing the presence of NADs and their role in repressing gene expression through the Polycomb repressive complex (PRC), as well as their correlation with differentiation [77, 79, 80]. Investigators confirm the interaction between nucleoli and NADs using fibrillin, nucleophosmin, or nucleolin as nucleolar markers. Interestingly, when mouse embryonic stem (ES) cells differentiated into neural progenitor cells (NPCs), a group of NADs migrated away the nucleolar periphery to the nucleus. Forty percent of the 550 genes belonging to these NAD group, which were linked to pathways involved in neuronal development and differentiation, are expressed, though their expression levels were varied greatly [79]. Ki-67 is not an authentic nucleolar marker. Nevertheless, our study demonstrated that Ki-67 disappearance due to MKI67 knockdown dispersed the condensed chromosomes from the nucleolar periphery to the nucleus, increasing the levels of 1,725 transcripts significantly (baseMean >15 and Pad <0.05). This gene group included the transcripts with the highest increases, i.e., those involved in neuronal differentiation, development, and activity (S5 Table). These results align with those observed in the mouse NPCs, suggesting that Ki-67 is causes or facilitates the NAD-nucleolus interaction and is responsible for neuronal differentiation, development, and maintenance. NPCs isolated from seven-day-old postnatal mice and grown in a dish formed spheres. Most cells in the spheres lost Ki-67 without differentiation treatment and lost it completely within a day of differentiation treatment [111]. This implies that the mouse NPCs obtained from ES cells by inducing differentiation, which were then used for NAD investigations, likely lost Ki-67 in many cells [79]. Consequently, the nucleolar peripheral NADs likely shifted slightly away from the nucleolar periphery within the nucleus in these cells. On the other hand, the PRC system is thought to silence cell-type-specific gene expression [77, 78, 112, 113]. Interestingly, many of the 1,725 transcripts increased by less than twofold, including WT1 transcripts, though some increased by as much as 15-fold. This high degree of variation in desilencing among the transcripts likely involves an additional factor in the silencing mechanism. Shafiq et al. demonstrated that the WT1 gene is regulated by the PRC alongside MTF2, which binds to CpG islands in the promoter region to maintain the silencing [114]. Their results imply that the low levels of WT1 desilencing resulting from Ki-67 depletion are due to MTF2 binding. This logic can be applied to the silencing of the 1,725 genes. Currently, the mechanism by which Ki-67 regulates the PRC is unknown. Additionally, we discovered that Ki-67 regulates translation by promoting the degradation of NEK7 transcripts. Therefore, Ki-67 regulates genomic transcript levels at either the transcription or translation level

While analyzing gene and protein expression, we found that increases in NEK7, POU4F2, PAX5, and SLC7A2 transcript levels, as well as changes in the transcript levels of glycolysis enzymes that increase ribose supply, can promote differentiation, development, and RNA and protein synthesis without rigorous proliferation [93–96]. Meanwhile, MDN1 coordinately regulates ribosome biogenesis and its energy supply under optimal conditions for proliferating cells. Taken together, these findings suggest that Ki-67, GNL2, and MDN1 collaborate to promote cell proliferation, including neural progenitor cell (NPC) self-renewal. They accomplish this by regulating the permeation of proteins and pre-60S particles through the nucleolar periphery, as well as the supply of ribosomes and energy. However, the disappearance of Ki-67 results in the expression of neuronal-cell-specific genes, leading to neuronal differentiation, development, and maintenance (Fig 11).

**Fig 11.**
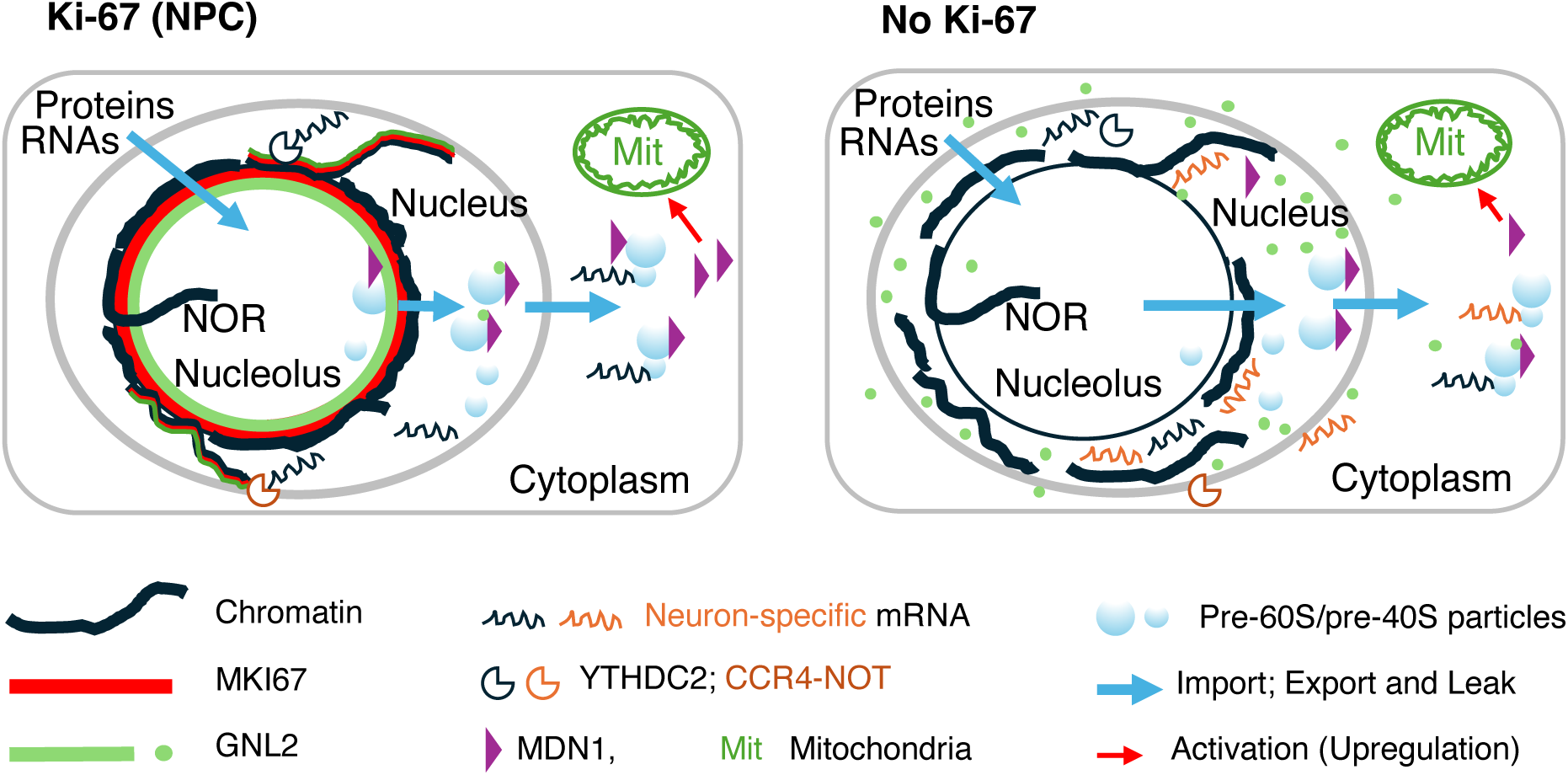
Illustration of changes in during neuronal differentiation induced by the absence of Ki-67. Ki-67 disappears during G0, which triggers changes for differentiation to neurons, and these changes persist in neurons. This figure and Fig 9D are complementary.

## Acknowledgements

We thank Dr. Danesh Moazed for his invaluable support and mentorship. We are grateful to Dr. Steven Gygi for access to the mass spectrometry facilities, and to Drs. Lingsheng Dong, Swapnil Suresh Parhad, William G. Rodriguez, and Harleen Saini for guidance on RNA-seq statistical analysis using the o2 high-performance computing platform. We appreciate Drs. Wenzhi Feng, Redi Metali, and Shuai Sun for training on the Applied Biosystems QuantStudio 7 Flex, and the CITE at Harvard Medical School for confocal microscopy access. We also thank the Biopolymers Facility Genomics Core (RRID:SCR_007175) for next-generation sequencing and the Harvard Chan Bioinformatics Core (RRID:SCR_025373)—especially Sergey Naumenko, Noor Sohail, and Shannan Ho Sui—for assistance with ChIP-seq data analysis. Their efforts were supported in part by the Harvard Medical School Foundry. We sincerely apologize to the authors of previous outstanding works for any incomplete citations. This was due in part to the limitations of our reference collection and constraints on available space within this paper.

## Author contributions

S.I. conceptualized the study, performed experiments, analyzed and visualized the data, and wrote the manuscript. J.A.P. conducted the mass spectrometry experiments, analyzed the data, wrote the corresponding methods section, and reviewed the manuscript.

## Funding

S.I. received support from the late Dr. Howard Green. J.A.P. was supported by an NIH grant (GM132129).

### Data availability

The plasmids utilized in this study related to the *GNL2* gene were deposited at Addgene. The ChIP-seq and RNA-Seq data were deposited in the Gene Expression Omnibus (GEO) database. The ChIP-seq accession number is GSE288519, and the RNA-seq accession numbers are GSE287864 and GSE287866. The MS data were deposited at Pride, and the project accession number is PXD070924.

### The websites utilized in this study

NoD (Nucleolar Localization Sequence Detector) at University of Dundee:

https://www.compbio.dundee.ac.uk/www-nod/

FastQC for ChIP-seq data:

https://www.bioinformatics.babraham.ac.uk/projects/fastqc/ developed by the Bioinformatics Group at the Babraham Institute. Fiji: http://imagej.org

Genome Browser

https://genome.ucsc.edu/cgi-bin/hgTracks?db=hg38&lastVirtModeType=default&lastVirtModeExtraState=&virtModeType=default&virtMode=0&nonVirtPosition=&position=chr7%3A155799529%2D155812871&hgsid=3431747607_msP9fAPiQF4IPocpT70KVYxAi9wq

Gene and Protein characteristics:

https://www.ncbi.nlm.nih.gov/datasets/gene/; https://www.uniprot.org/uniprotkb

Cytoscape: https://cytoscape.org/

Metascape: https://metascape.org/gp/index.html#/main/step1

AlphaFold 3: https://alphafoldserver.com/welcome

## Supporting figure captions

**S1 Fig.**
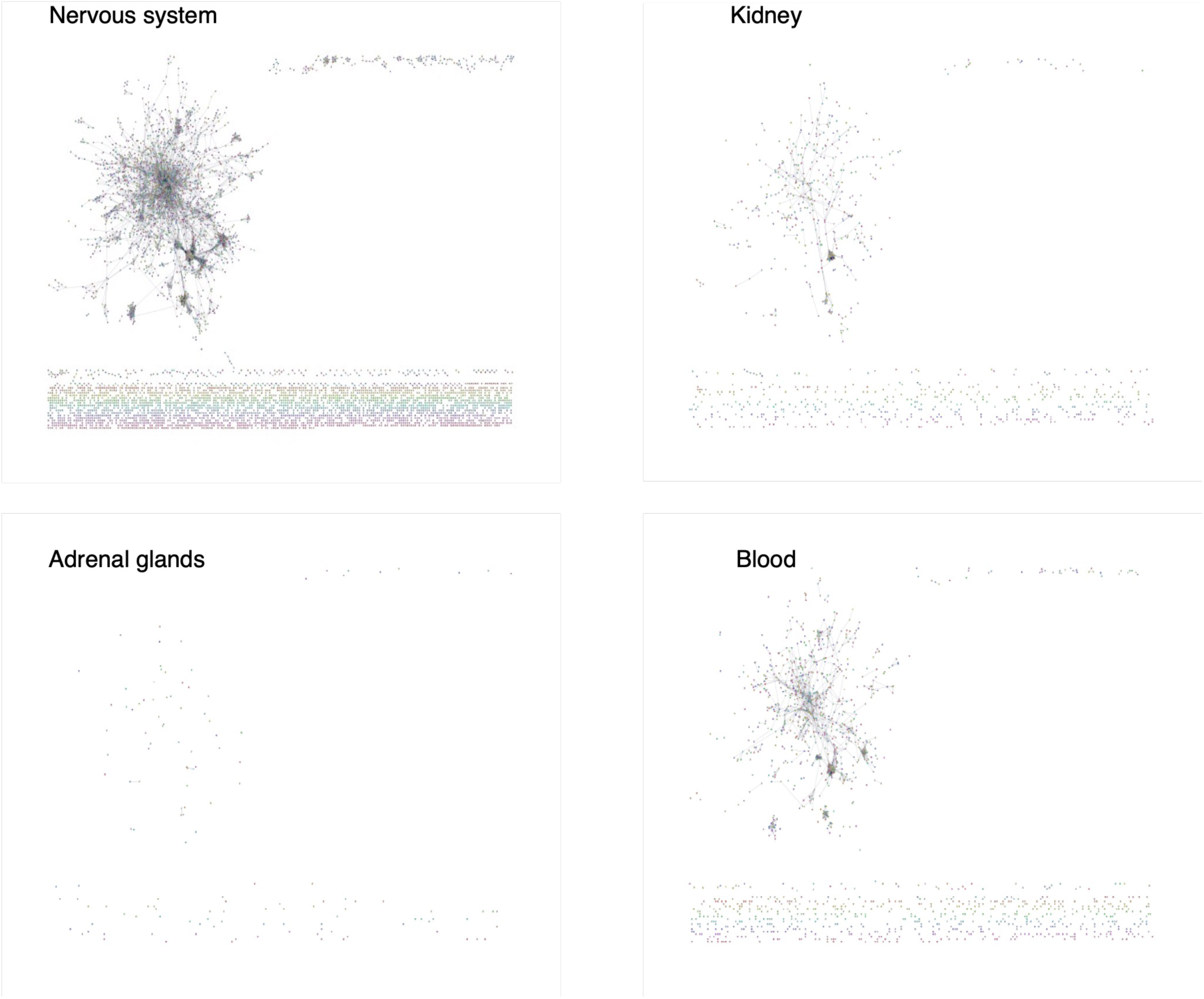
Nuclear proteins expressed in HEK293T cells exhibited the greatest similarity to those expressed in the human nervous system. This similarity was evaluated using the second-most-severe tissue specificity level out of five.

**S2 Fig.**
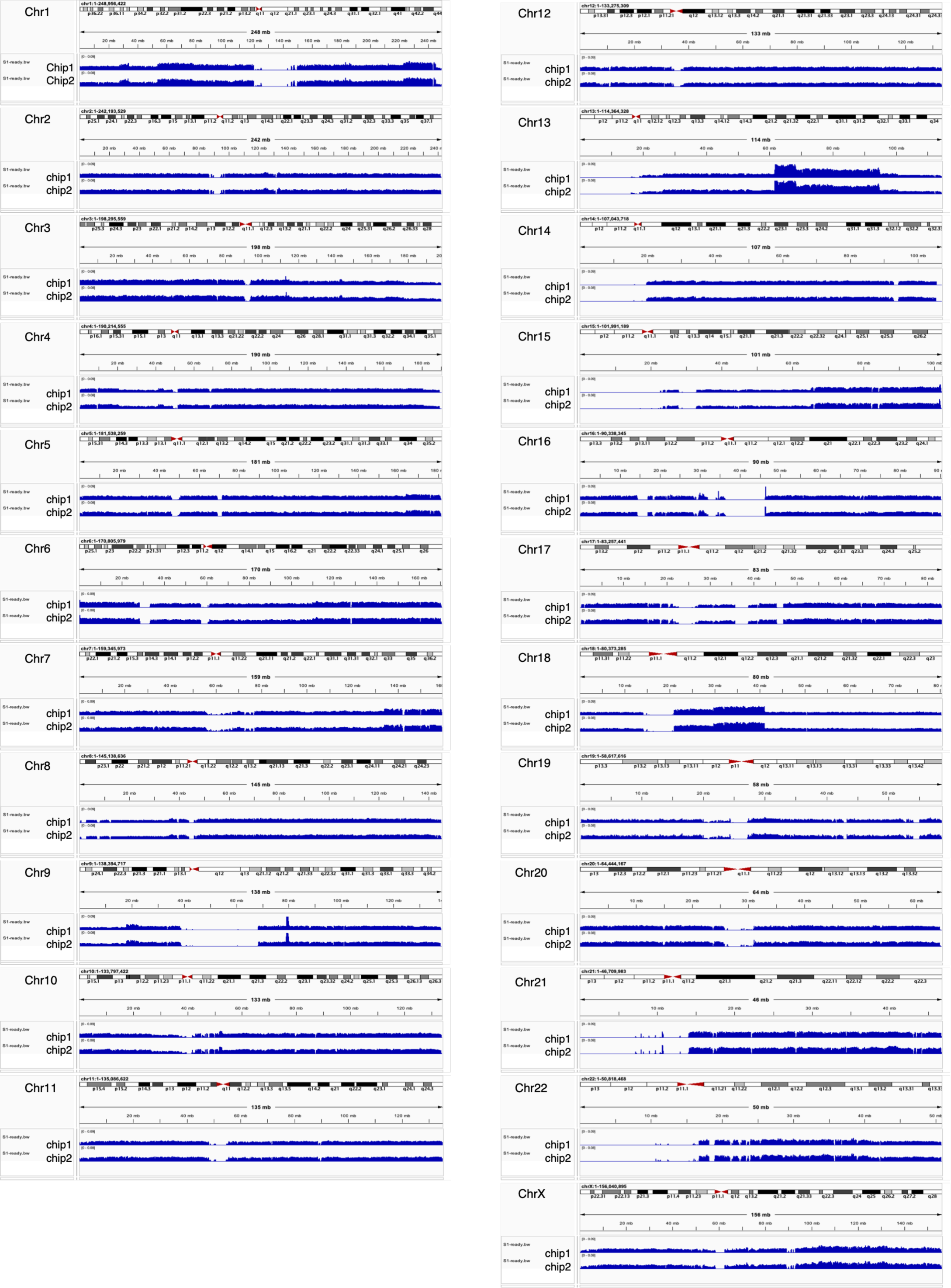
BigWig presentation of Ki-67 ChIP-seq results on all chromosomes. Consistent Ki-67 binding between chip1 (the initial ChIP-seq) and chip2 (a second ChIP-seq using an independently prepared sample) revealed the validity of the data.

**S3 Fig.**
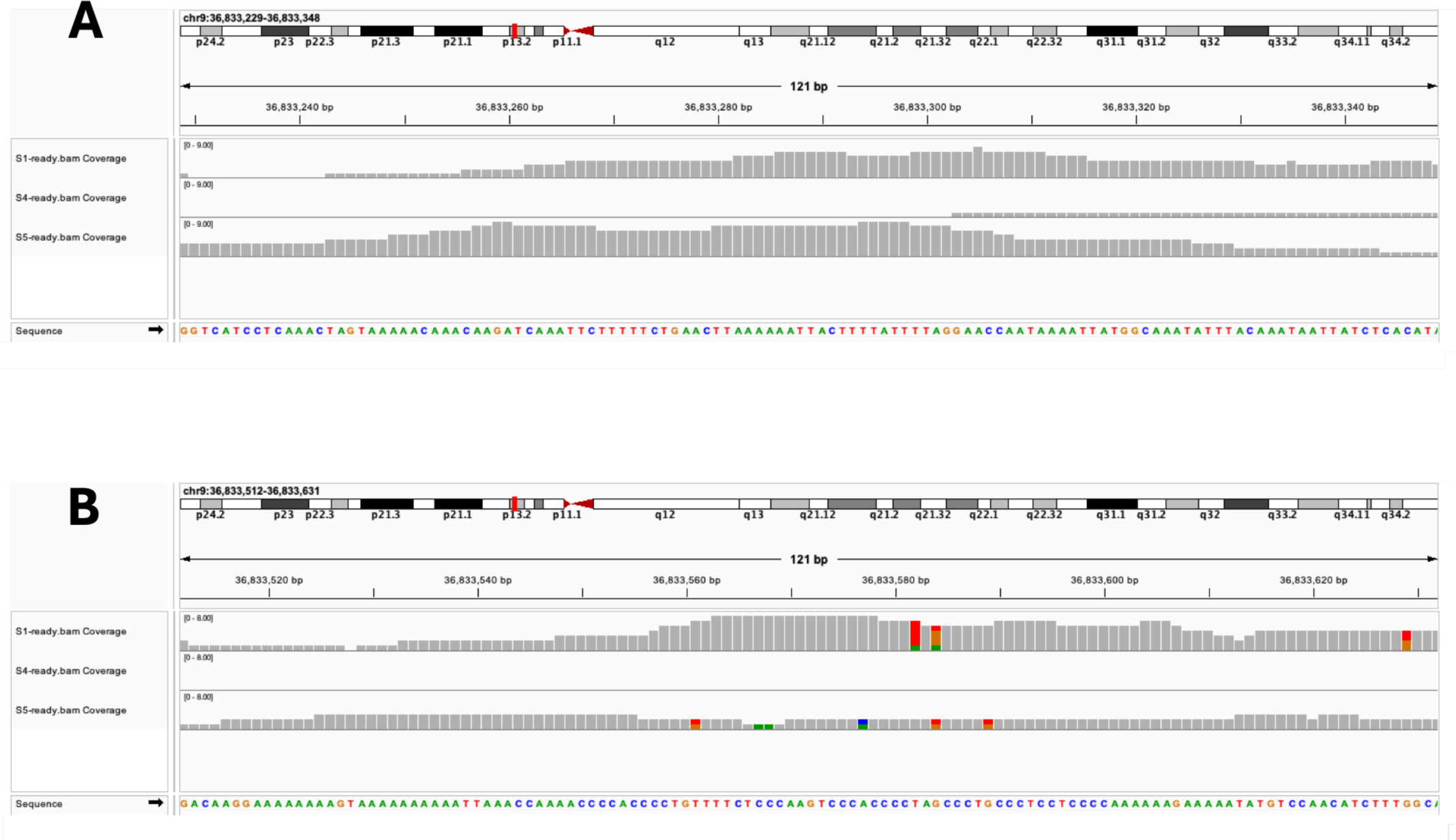
Ki-67 binding to AT-rich nucleotide sequences and GC-rich nucleotide sequences. **A**. A nucleotide-level view of Ki-67 binding to an AT-rich sequence presented by BAM tracks. The first, second, and third tracks show Ki-67 ChIP-seq reads, the control, and the input, respectively. **B**. A nucleotide-level view of Ki-67 binding to a GC-rich sequence presented similarly to (A).

**S4 Fig.**
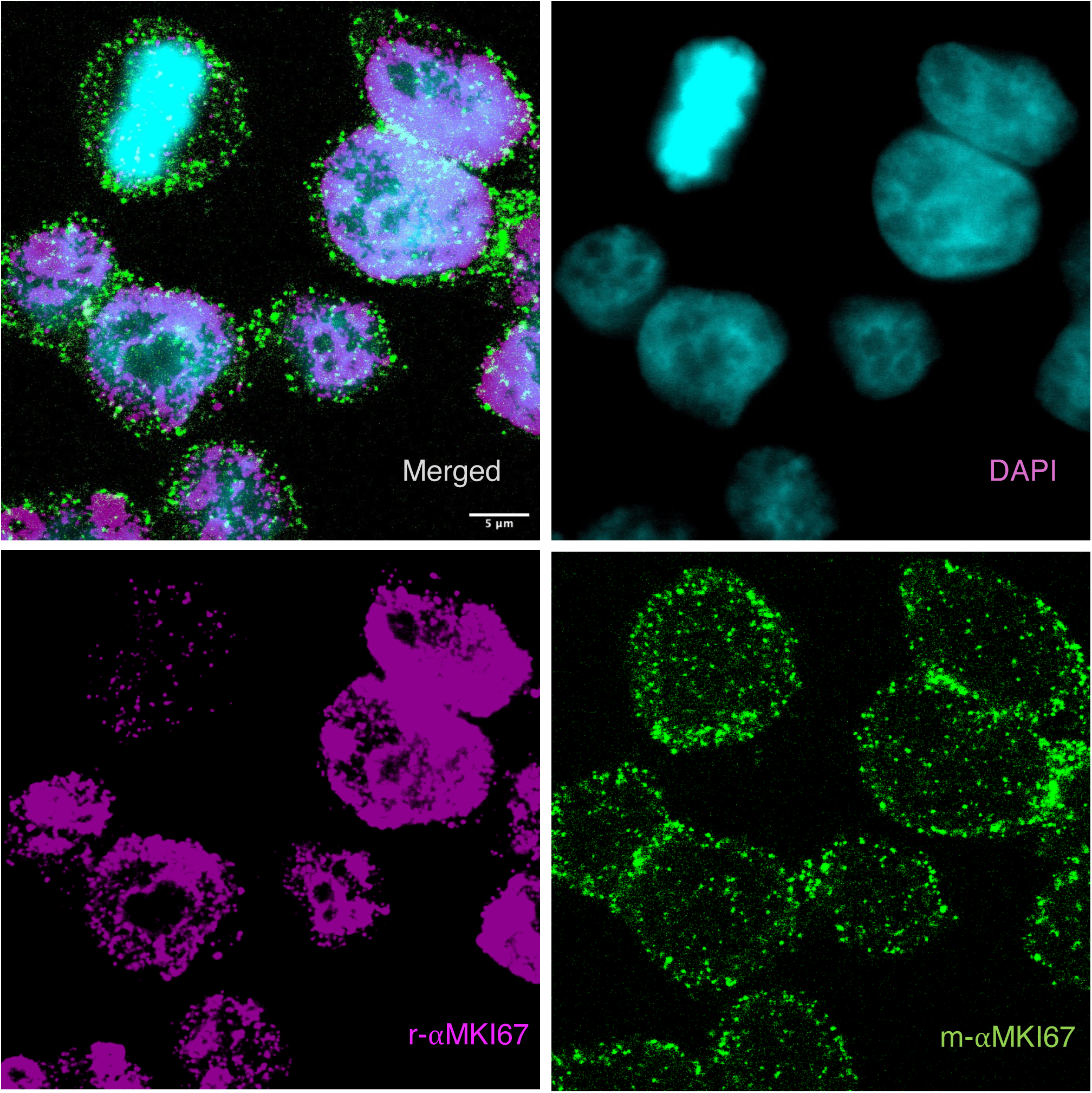
The rabbit monoclonal anti-Ki-67 antibody (r-αMKI67) that we used throughout this study detected Ki-67 at a level of 7% the intensity of interphase Ki-67. This was determined by dividing the signal intensity ratio of mitotic cells by that of interphase cells. We calculated the signal intensity ratio of mitotic cells by dividing the signals detected with r-αMKI67 by the signals detected with the mouse monoclonal antibody m-αMKI67. The signal intensity ratio of interphase cells was calculated similarly. The proportion of mitotic cells was 6% of the total number of cells in the randomly proliferating cultures. Therefore, we estimated that the noise value driven from mitotic cells was 0.42 % (0.07 x 0.06 x 100), which had a negligible effect on the results. Images for MKI67 were obtained with overexposure to visualize even weak signals.

**S5 Fig.**
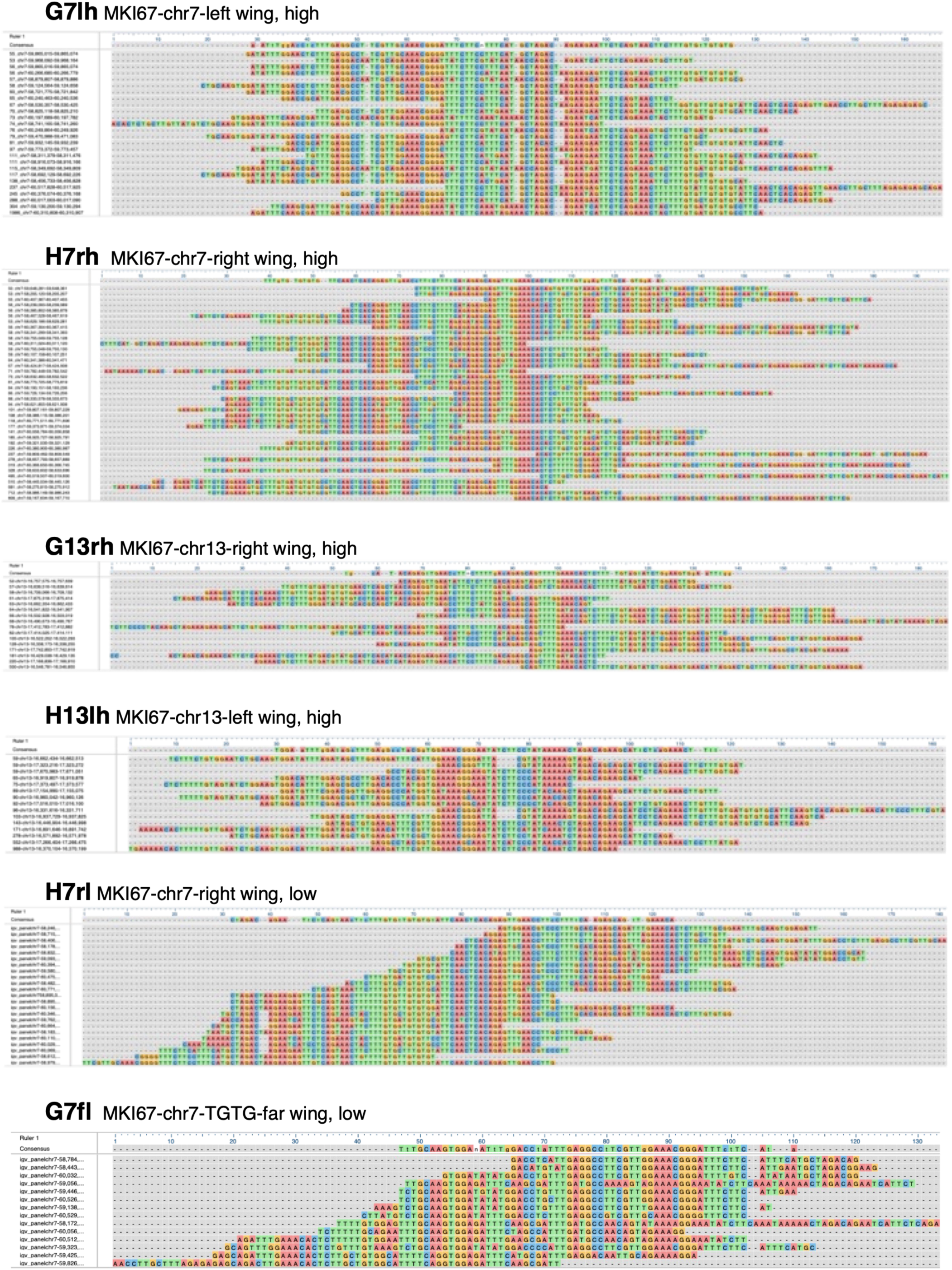
Alignment of the Ki-67 binding region sequences in chromosomes 7 and 13. This procedure was performed separately for the high- and low-binding groups on chr7. The same procedure was performed for chr13. The total number of reads at the Ki-67 binding site was counted and placed in ascending order on the line farthest left line. The corresponding nucleotide sequences for each count were placed to the right. The consensus sequences for these alignments are shown above.

**S6 Fig.**
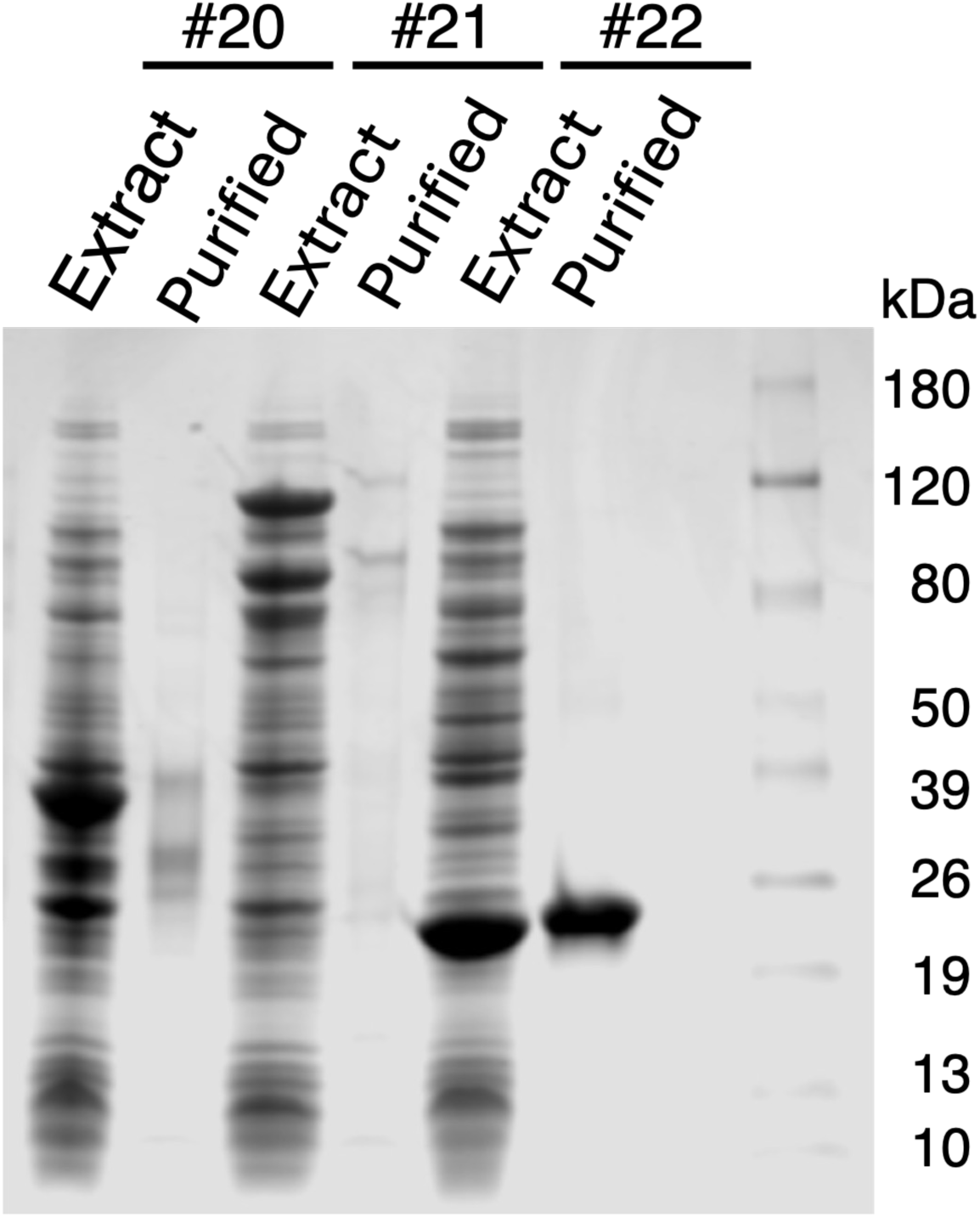
Affinity purification of probes. The probes, which were expressed in E. coli, were bound to Ni-NTA magnetic agarose, washed, and finally eluted. The probes obtained through these steps (#20 and #21) were pure, though they contained cleaved fragments. Probe #22 consists only of the EGFP-6xHis tag.

**S7 Fig.**
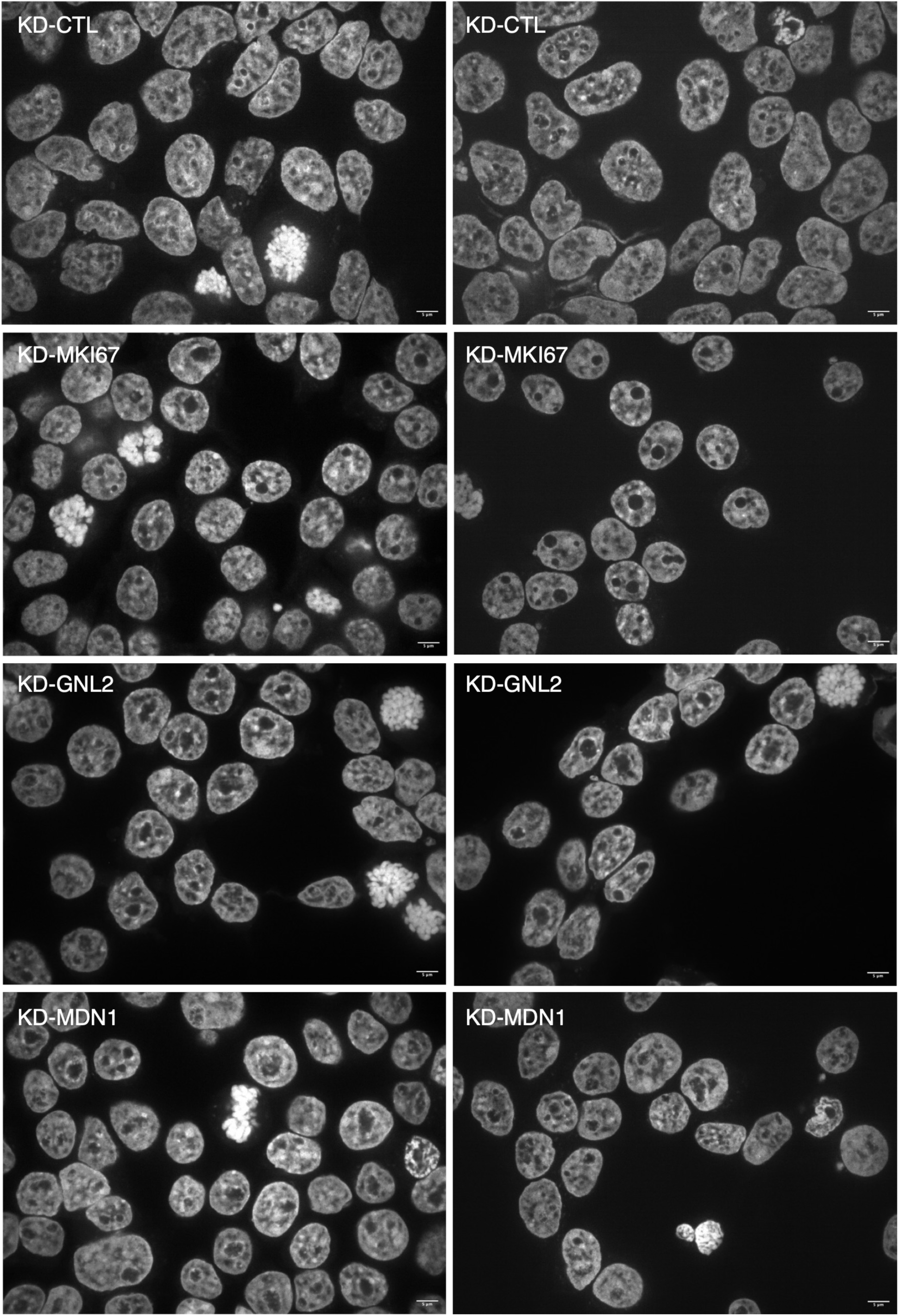
DAPI-stained nuclei of *MKI67-*, *GNL2-*, and *MDN1*-knockdown cells, along with the control cells.

**S8 Fig.**
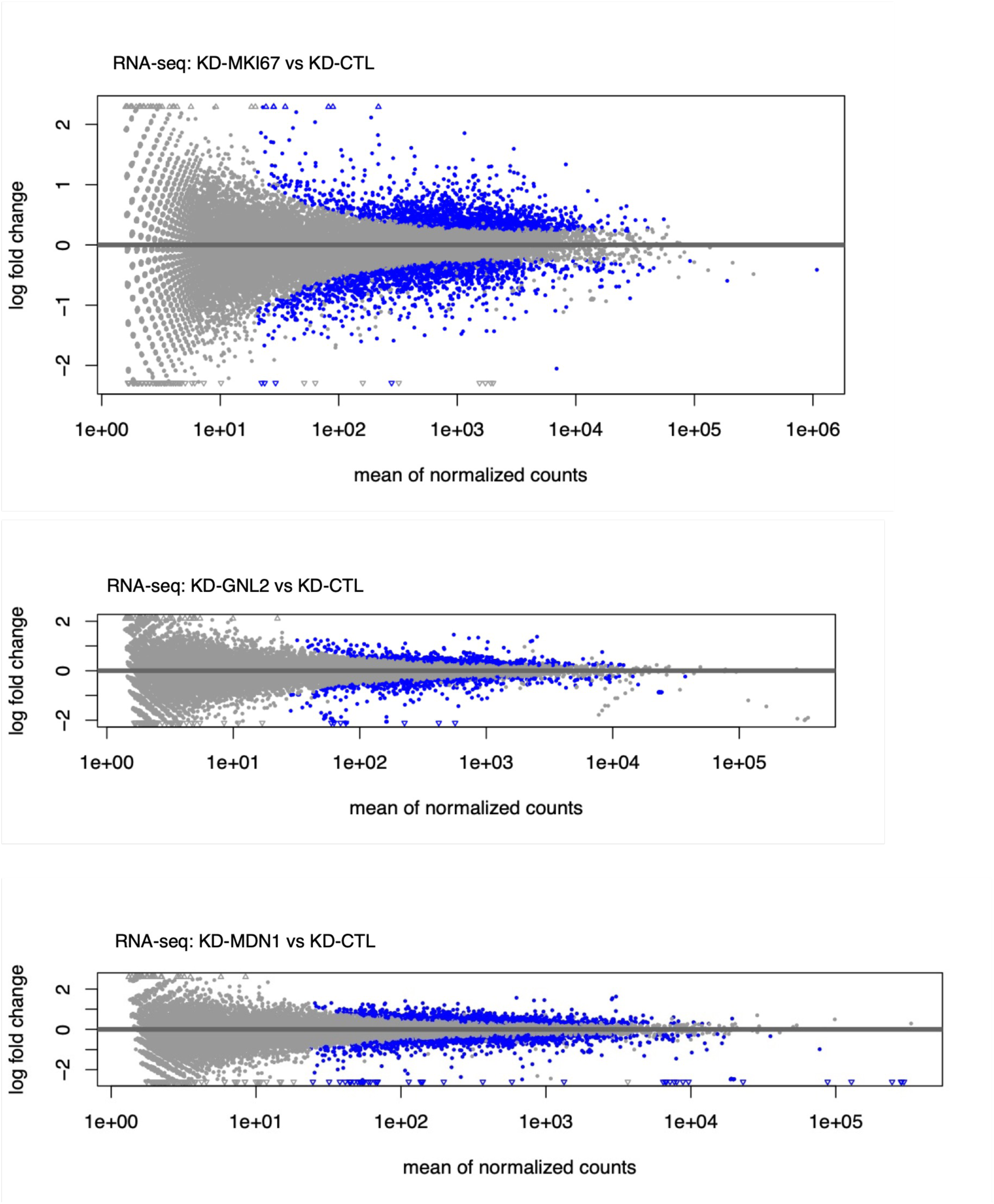
Mplot: genomic transcript level changes through *MKI67*, *GNL2*, and *MDN1* knockdown from control levels.

**S9 Fig.**
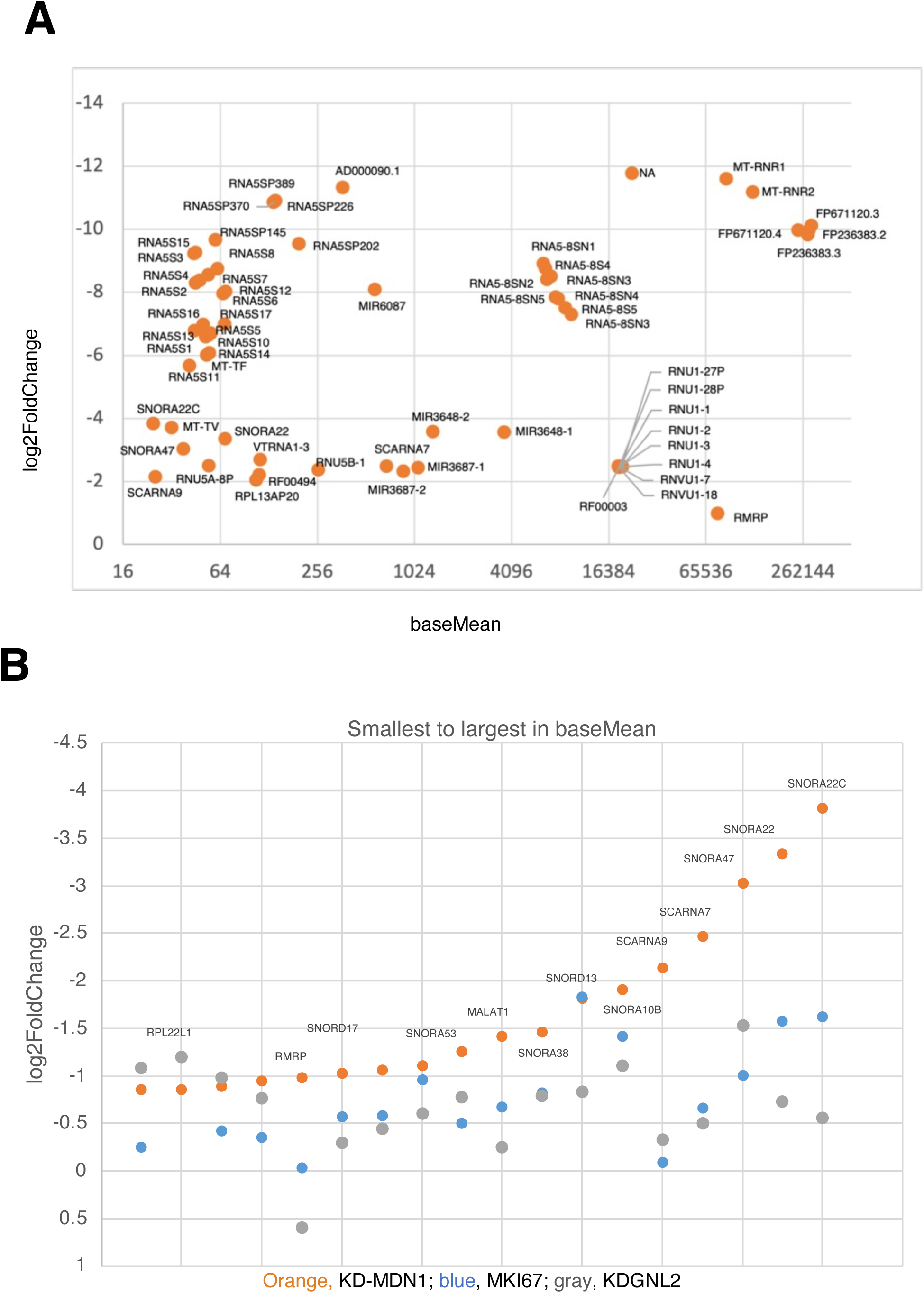
**A**. Mplot of transcripts that decreased markedly through *MDN1* knockdown. **B**. *MDN1* knockdown decreased transcript levels more severely than *MKI67* and *GNL2* knockdowns.

**S10 Fig.**
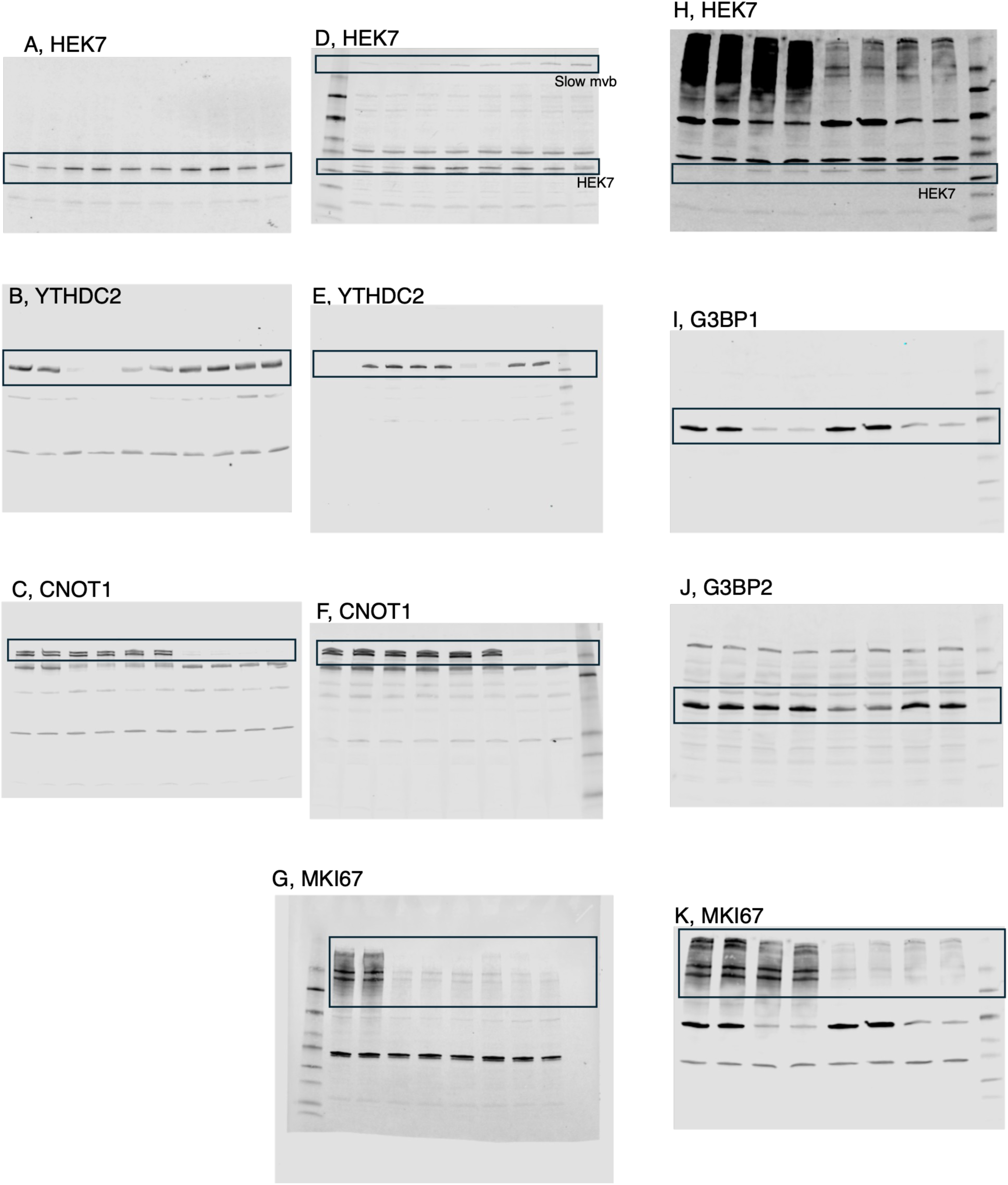
**A-K**. Western blot originals. The framed selections were used to create Fig 7. Some of the images exhibited noise bands. This is attributed to the use of two primary antibodies in the Western blotting process, or to a second Western blot that targets a different protein after the initial identification.

**S11 Fig.**
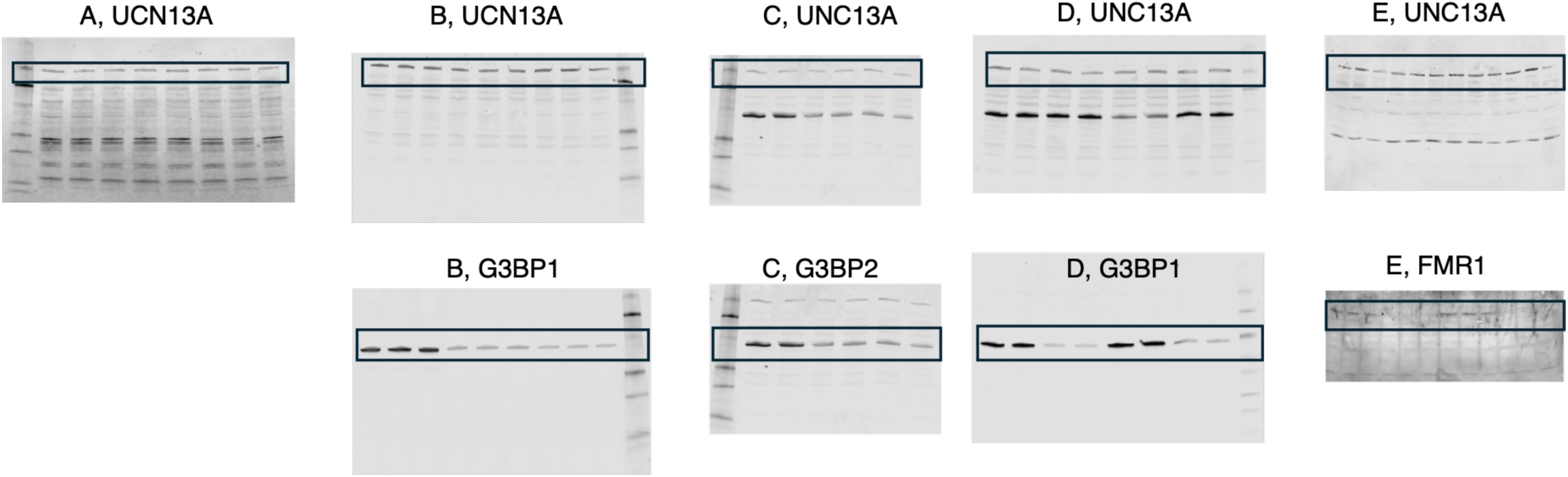
**A-E**. Western blot originals. The framed selections were used to create Fig 8. The top images show the levels of UNC13A resulting from various knockdowns. The images below confirm the knockdown. Some of the images exhibited noise bands. This is attributed to the use of two primary antibodies in the Western blotting process, or to a second Western blot that targets a different protein after the initial identification.

**S12 Fig.**
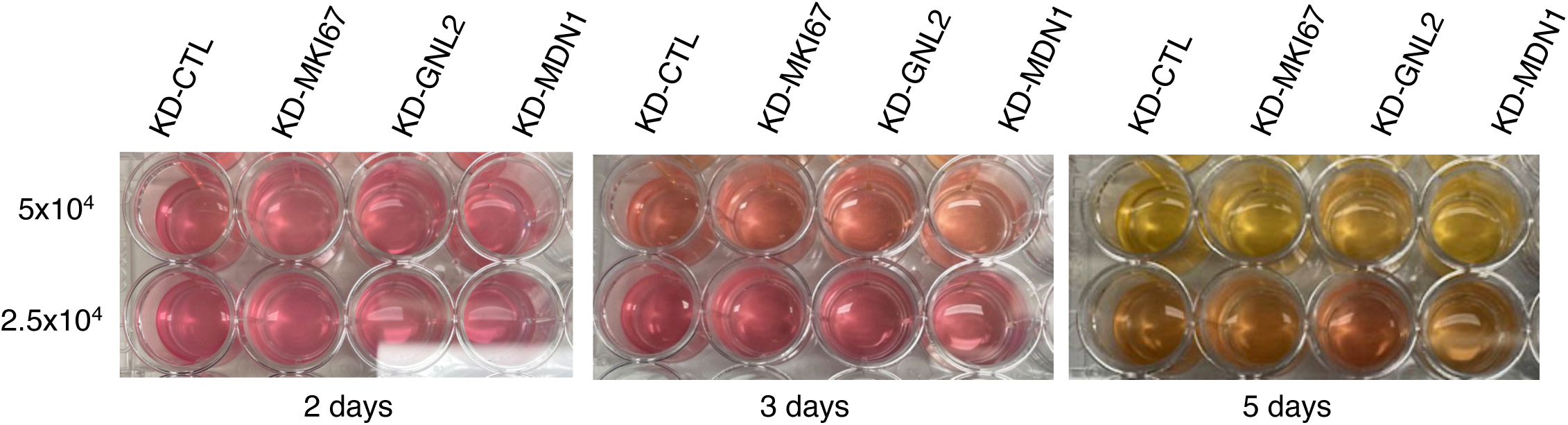
Acidification observed in cultures knocked down for the *MKI67*, *GNL2*, or *MDN1* genes, as well as in the negative control culture. The three genes were knocked down in separate wells of a 24-well plate. The number of cells per well was adjusted to either 5x10⁴ or 2.5x10⁴ for each knockdown. The total volume in each well was 1.12 ml. The plate was then incubated for five days.

**S13 Fig.**
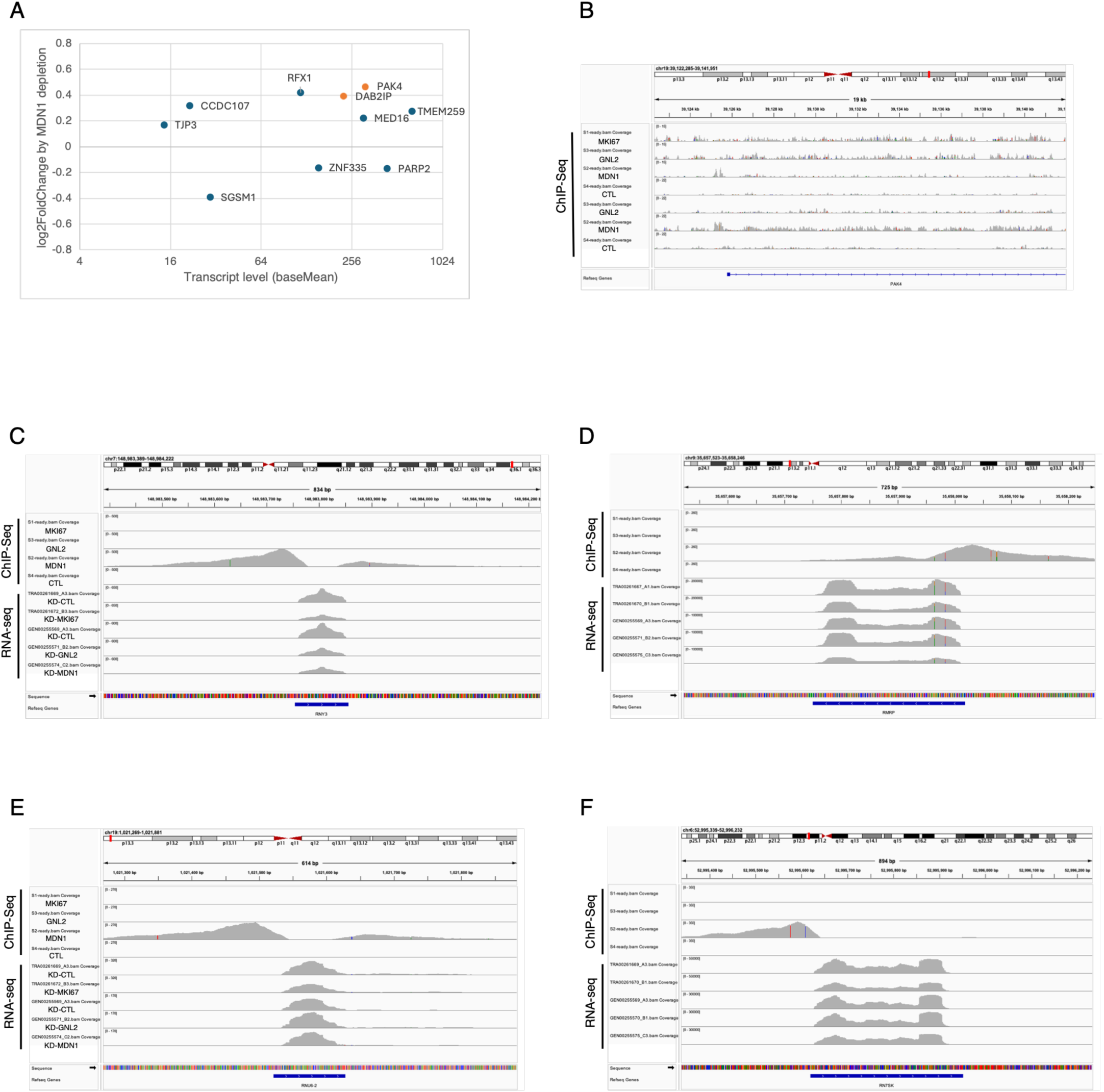
**A.** Mplot of transcripts from genes whose promoter regions were bound by MDN1. **B.** MDN1 binding to the *PAK4* promoter region. **C**. MDN1 binding to the *RNY3* promoter region and *RNY3* transcript levels. **D**. MDN1 binding to the *RMRP* gene and *RMRP* transcript levels. **E**. MDN1 binding to *RNU6-2* and *RNU6-2* transcript levels. **F**. MDN1 binding to *RN7SK* and *RN7SK* transcript levels.

**S14 Fig.**
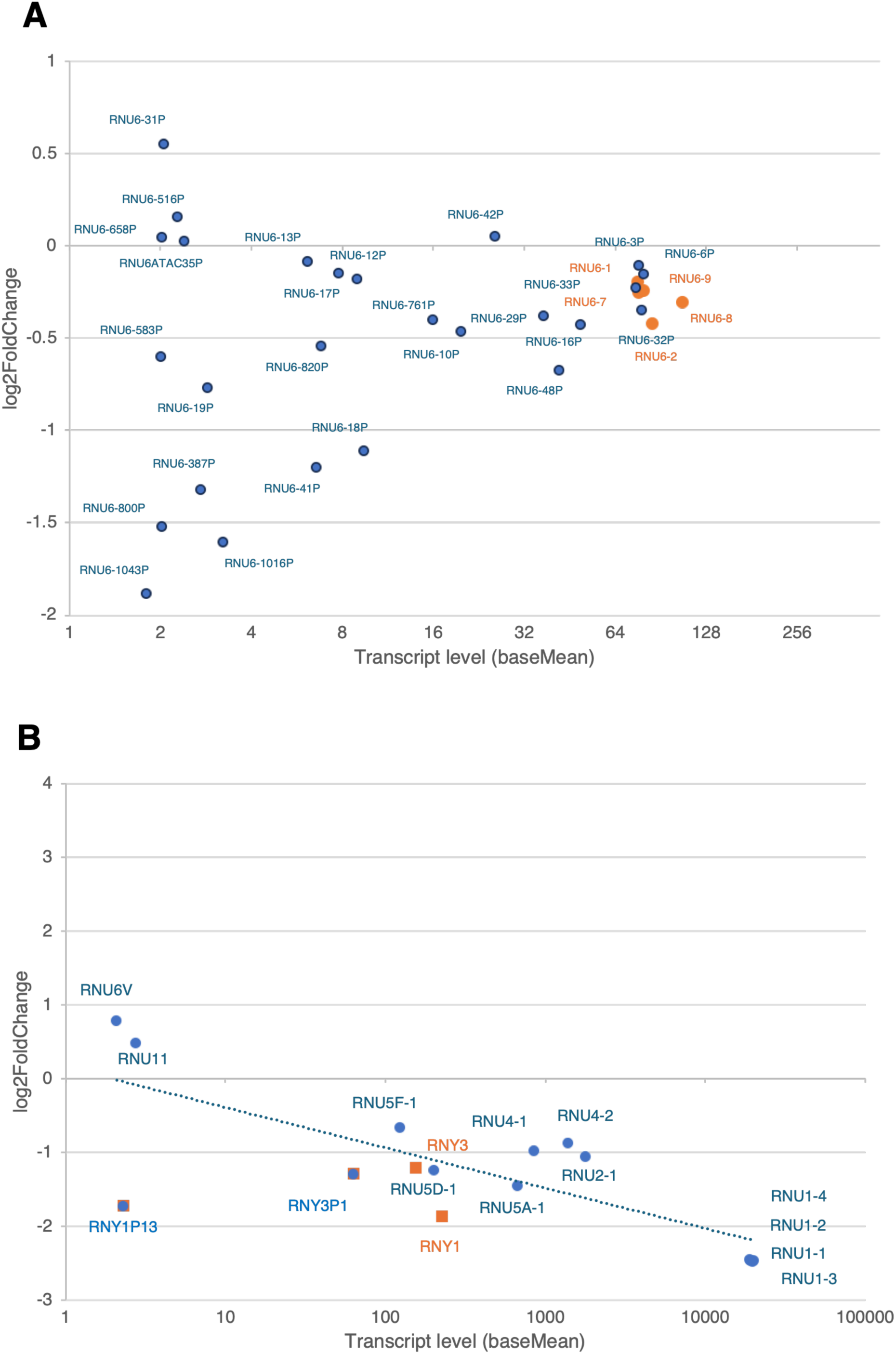
Mplot illustrates the changes in RNU6 transcript levels resulting from KD-MDN1. These changes (orange) were similar to those of their pseudogene transcripts, to which MDN1 did not bind. B. Mplot shows the change in RNY transcript levels resulting from KD-MDN1. The changes in the RNY transcript levels resulting from MDN1 knockdown (orange) were similar to the changes in the pseudogene transcript levels.

