## Supplementary figures and images for "The roles of MKI67, GNL2, and MDN1 in Ribosome biogenesis and Transcriptome regulation in the Neuronal Lineage cell line HEK293T"

### S1 Table.docx

S1A Table

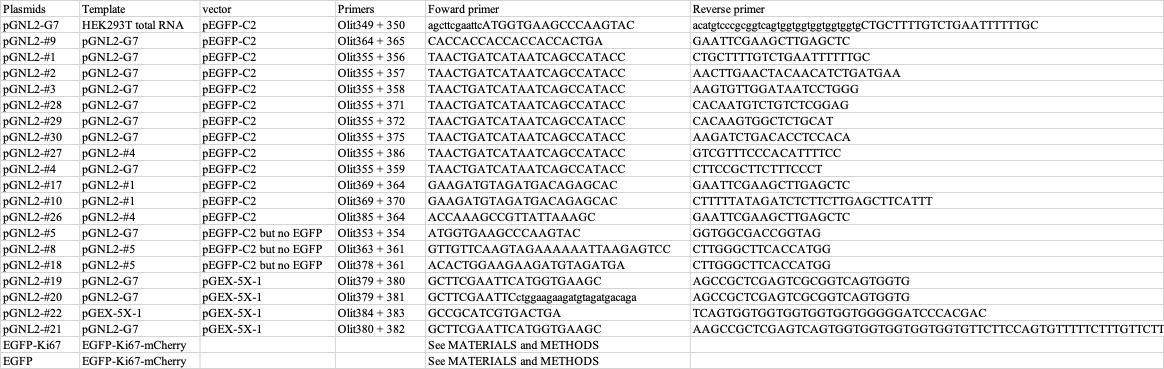

S1B Table

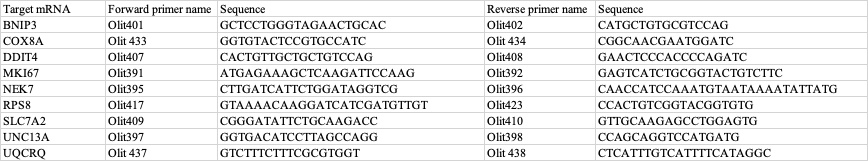

### S2 Table.docx

**S2 Table**

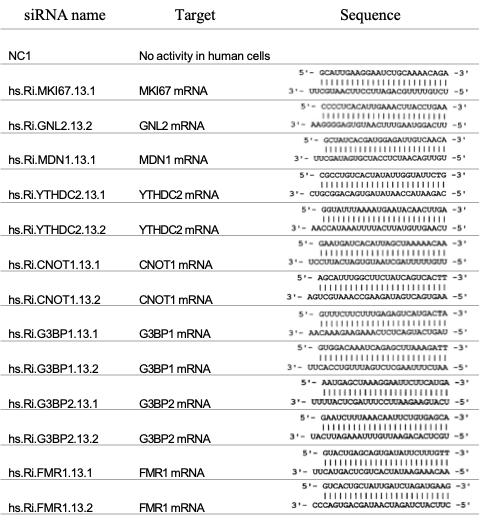
