## Supporting Table 3 for "The roles of MKI67, GNL2, and MDN1 in Ribosome biogenesis and Transcriptome regulation in the Neuronal Lineage cell line HEK293T": S3 Table.docx

| Target | Antibody | Host and Type | Concentration | (µg/ ml) | | Cat# | Vendor |
| --- | --- | --- | --- | --- | --- | --- | --- |
| - | - | - | IF | WB | IP | - | - |
| CNOT1 | CNOT1 (D5M1K) | r-Monoclonal |  | 0.1 |  | 44613 | Cell Signaling |
| FMR1 | Anti-FMR1 (1F1) | m-Monoclonal |  | 1 |  | MABN2453 | MILLIPORE |
| G3BP1 | G3BP1 (E9G1M) XP | r-Monoclonal |  | 0.075 |  | 61559S | Cell Signaling |
| G3BP2 |  |  |  | 0.031 |  | 31799S | Cell Signaling |
| GFP | GFP antibody, purified | r-Polyclonal |  | 0.25 | 5-10 | SP3005P | ORIGENE |
| GNL2 | GNL2 (B-8) | m-Monoclonal | 2.5 | 1 |  | sc-514050 | SANT CRUZ |
| GNL2 | GNL2 antibody | r-IgG |  |  | ~5 | A305-155A-M | BETHYL |
| 6x His | Tetra.His Antibody | m-IgG1 | 1 |  |  | 34670 | QIAGEN |
| 6x His | U571WHA040-1 | m-Polyclonal |  | 0.1 | ~10 | A00186-100 | GenScript |
| MDN1 | MDN1 antibody | r-Polyclonal | 0.7 | 0.25 | ~5 | PA556225 | Thermo Fisher |
| MKI67 | Human Ki-67/MKI67 antibody | r-Monoclonal | 1 | 0.25 | ~5 | MAB7617-SP | R&D |
| MKI67 | Ki-67 (Ki67) | m-Monoclonal | 8 |  |  | sc-23900 | SANT CRUZ |
| NEK7 | NEK7 (C34C3) | r-Monoclonal |  | 0.04 |  | 3057S | Cell Signaling |
| PELP1 | PELP1 (D5Q4W) | r-Monoclonal | 0.94 |  |  |  | Cell Signaling |
| UNC13A |  | r-IgG |  | 0.2 |  | 55053-1-AP | proteintech |
| YTHDC2 | YTHDC2 | r-IgG |  | 0.2 |  | 27779-1-AP | proteintech |
| mouse IgGs, IgM, IgA | IRDye 680RD Goat anti-Mouse IgG | goat-Polyclonal |  | 0.025 |  | 926-689070 | LI-COR |
| rabbit IgG | IRDye 800 Affinity Purified Goat anti-Rabbit IgG | goat-Polyclonal |  | 0.05 |  | 611-132-122 | ROCKLAND |
| mouse IgG (H+L) | Goat anti-Mouse Highly Cross-Absorbed, Alexa Fluor 488 | goat-Polyclonal | 1 |  |  | A11029 | Thermo Fisher |
| rabbit IgG (H+L) | Goat anti-Rabbit Highly Cross-Absorbed, Alexa Fluor 555 | goat-Polyclonal | 1 |  |  | A21429 | Thermo Fisher |
